# Typical and disrupted brain circuitry for conscious awareness in full-term and preterm infants

**DOI:** 10.1101/2021.07.19.452937

**Authors:** Huiqing Hu, Rhodri Cusack, Lorina Naci

## Abstract

One of the great frontiers of consciousness science is understanding how early consciousness arises in the development of the human infant. The reciprocal relationship between the default mode network (DMN) and frontoparietal networks — the dorsal attention network (DAN) and executive control network (ECN) — is thought to facilitate integration of information across the brain and its availability for conscious access to a wide set of mental operations. It remains unknown whether the brain mechanism of conscious awareness is instated in infants from birth. To address this gap, we asked what the impact of prematurity and neonate age is on the development the default mode and fronto-parietal networks, and of their reciprocal relationship. To address these questions, we used the Developing Human Connectome Project (dHCP), a unique Open Science project which provides a large sample of neonatal functional Magnetic Resonance Imaging (fMRI) data with high temporal and spatial resolution. Resting state fMRI data for full-term neonates (N = 282, age 41.2 w ± 12 d), and preterm neonates scanned at term-equivalent age (TEA) (N = 73, 40.9 w ± 14.5 d), or before TEA (N = 73, 34.6 w ± 13.4 d) were obtained from the dHCP, and for a reference adult group (N = 176, 22 – 36 years), from the Human Connectome Project. For the first time, we show that the reciprocal relationship between the DMN and DAN was present at full-term birth or TEA. Although different from the adult networks, the DMN, DAN and ECN were present as distinct networks at full-term birth or TEA, but premature birth disrupted network development. By contrast, neonates before TEA showed dramatic underdevelopment of high-order networks. Only the DAN was present as a distinct network and the reciprocal network relationship was not yet formed. Our results suggest that, at full-term birth or by term-equivalent age, infants possess key features of the neural circuitry that enables integration of information across diverse sensory and high-order functional modules, giving rise to conscious access. Conversely, they suggest that this brain infrastructure is not present before infants reach term-equivalent age. These findings improve understanding of the ontogeny of high-order network dynamics that support conscious awareness, and of their disruption by premature birth.

## Introduction

It remains unknown whether conscious awareness is present in newborn infants and whether its development is affected by premature birth. In healthy human adults, consciousness is clinically defined and measured by examining two distinct dimensions: arousal and awareness (Laureys et al., 2004). ‘Arousal’ is measured by assessing spontaneous eye opening, sleep-wake cycles and other systemic fluctuations in the ability to engage with the environment. ‘Awareness’ is assessed by examining the ability to wilfully to respond to commands behaviourally and/or through language, as well as the ability to report on mental states pertaining to oneself or the environment (Naci et al., 2014; Mehling et al., 2009; Clare et al., 2005). There is no question that newly born infants, or neonates, have arousal, e.g., they cry and have sleep-wake cycles. However, the extent to which neonates can consciously process information about themselves and their environment remains unknown. Highly relevant to understanding neonate awareness is Damasio’s (2000) distinction between a ‘core awareness’, or a basic integrated experience of the current moment, and an ‘extended awareness’, made possible by the accumulation of autobiographical memories that allow creation of an internal world and projection beyond the present. This is echoed in the distinction between a ‘minimal’ and ‘longitudinal’ self, proposed by others. The ‘minimal’ self is immediate (Gallagher, 2000) and comprises awareness of body boundaries and position, facial features and body size, visceral states, agency, mental states and online behaviour (Sturm et al., 2017), whereas the ‘longitudinal’ self, requires the presence of episodic autobiographical memory (i.e., memories of past events) and semantic self-knowledge (i.e., knowledge of one’s own traits) and is extended across time (Seeley and Sturm, 2006).

Although studies on neonate awareness are scarce, a few suggest ‘minimal’ awareness from birth. Newborns perform some forms of stimulus discrimination early after birth, including distinguishing their body (12 to 103 hours after birth; Filippetti et al., 2013), and their own cry from those of other newborns’ (within one day after birth; Martin et al., 1982; Simner et al., 1971), their mother’s voice from a stranger’s (within 12 hours – 3 days after birth; Ockleford et al., 1988; Querleu et al., 1984; DeCasper et al., 1980), and discriminating facial expressions of happiness from disgust (2 days after birth; Addabbo et al., 2018). Although prima facia, these studies suggest ‘minimal’ awareness in neonates, these behaviours could be due to certain stimuli being primed in the early days of life or even in the womb, or due to the physical properties of the stimuli themselves. For example, the mother’s voice is very familiar to the neonate, so any preferential responses could be due to familiarity rather than understanding the meaning and significance of the mother figure. Similarly, the response to expressions of disgust, could be a pre-conscious reaction to aversive stimuli (Ruffman et al., 2019). Critically, the lack of language and the very limited motor function preclude self-report or behavioural responses, and, thus, prevent the assessment of infant awareness from the first days of life. To circumvent these limitations, in present study we followed a different strategy.

We asked a foundational question to understanding the *capacity* for conscious experiences, that of whether or not the brain mechanisms of conscious awareness are instated in neonates. We focused in particular on the development of the fronto-parietal and default mode (DMN) networks, of their reciprocal relationship, and how they are impacted by premature birth and neonate age. Prominent theories of consciousness, including the Global Neuronal Workspace Theory (Mashour et al., 2020; Dehaene et al., 2011) and the Integrated Information Theory (Tononi, 2004) converge on the principle that consciousness requires integration of information from discrete but interconnected modules across the brain. Adult functional neuroimaging studies have identified the fronto-parietal and DMN networks as two such distinct cortical systems that support consciousness and play complementary roles in information integration. Myriad neuro-psychological and neuroscientific studies show that fronto-parietal regions — comprising the dorsal attention (DAN) and executive control (ECN) networks — are critical for stimulus-driven high-order cognition (Ptak, 2012; Woolgar et al., 2010; Duncan et al., 2010; Elliott, 2003; Shallice, 1988), and recent work suggests they facilitate awareness of external stimuli (Huang et al., 2020; Demertzi et al., 2013; Vanhaudenhuyse et al., 2011). By contrast, the DMN has been primarily implicated in self-referential processing (Qin and Northoff, 2011; Andrews-Hanna et al., 2010; Schneider et al., 2008; Beer, 2007; Buckner et al., 2007; D’Argembeau et al., 2005; Wicker et al., 2003; Gusnard et al., 2001) and self-awareness (Demertzi et al., 2013; Vanhaudenhuyse et al., 2011), and more recently in context processing (Smith et al., 2018; Vatansever et al., 2018; Margulies et al., 2016). Importantly, the fronto-parietal network and DMN share a reciprocal relationship, where they are not simultaneously active, i.e., are anticorrelated, or exhibit low correlation of functional time-courses relative to other brain network pairings. This relationship is abolished when consciousness is extinguished, irrespective of condition, e.g., whether during deep anaesthesia under various pharmacological manipulations or after severe brain injury (Huang et al., 2020; Haugg et al., 2018; Bonhomme at el., 2012), suggesting that it tracks the presence of conscious awareness.

Whether the DMN, DAN and ECN and their reciprocal relationship are developed by birth and whether they are affected by prematurity, remain poorly understood. To assess the literature, we conducted a literature search with key words, ‘functional network’ or ‘functional connectivity’, ‘infant’ or ‘newborn’ or ‘neonatal’, and “fMRI” that resulted in 19 neonate studies summarized in Table S1. While the three primary sensory and motor networks were consistently reported in neonates (Cui et al., 2017; Gao et al., 2015a, 2015b; Doria et al., 2010; Fransson et al., 2009; Fransson et al., 2007), findings were inconsistent on the presence of high-order networks, including the DMN, DAN, ECN. Some resting state functional magnetic resonance imaging (rs-fMRI) studies (Gao et al., 2015a, 2009; Smyser et al., 2010; Fransson et al., 2007) found no evidence for the presence of these networks until the end of the first year. Fransson et al. (2007) found that the DMN in preterm neonates was fragmented into an anterior and posterior part. Similarly, Gao et al., (2009) reported that although the two main hubs of DMN (i.e., the ventral/dorsal medial prefrontal cortex and posterior cingulate/retrosplenial cortex) were consistently observed in 2-week-olds, 1-year-old and 2-year-olds, other aspects of the DMN (i.e., the inferior parietal lobule, lateral temporal cortex, and hippocampus regions) were not found in 2-week-olds. A similar pattern was also reported for the DAN and ECN (Gao et al., 2015b; Fransson et al., 2009). By contrast, other rs-fMRI studies (Linke et al., 2018; He et al., 2015, 2016; Doria et al., 2010) support the idea that these networks have already emerged in neonates. For instance, Doria et al. (2010) found that both primary and high-order networks were present in full-term and preterm neonates scanned at term-equivalent age (TEA: 37 – 42 weeks of postmenstrual age). He et al. (2015) detected a fronto-parietal network, comprising the frontal gyrus and inferior parietal cortex, and a second one, comprising the anterior cingulate cortex, medial prefrontal cortex, superior/middle frontal gyrus, in preterm neonates. Linke et al. (2018) found that both the ECN and DMN were present even in neonates with perinatal brain injuries, both full-term and preterm neonates scanned at TEA.

Several factors may contribute to these divergent results. Due to methodological and technical challenges, the incipient field of infant neuroimaging has, to date, not adhered to unified testing protocols. The aforementioned studies employ vastly different sample sizes (e.g., ranging from N = 11 to 143), different MR field strengths (e.g., 1.5T vs 3T) yielding different spatial and temporal resolutions (Weiskopf et al., 2006; Triantafyllou et al., 2005), different motion artifact control methods, and lack age-specific structural brain templates for neonates. The multitude of different methodologies across previous studies render it impossible to conclude, in light of inconsistent results, whether the DMN, DAN and ECN are already present at birth or not. Moreover, to the best of our knowledge, only one study to date (Gao et al., 2013) has investigated whether the reciprocal relationship between these three high-order networks is developed in early infancy. Gao et al. (2013) reported that the anticorrelated interaction between the DMN and DAN was absent at birth, but became apparent at one year of age. However, this study had a relatively small number of full-term neonates (N = 51) and no preterm neonates, which may have reduced the power to detect effects of interest. To address aforementioned limitations of previous studies, we used data from the open-source Developing Human Connectome Project (dHCP). The dHCP conferred several advantages, including a robust sample size (N = 282), 3T MRI, multiband echo-planar imaging that significantly improves temporal resolution and signal-to-noise (Zhang et al., 2019), registration to more accurate week-to-week neonate structural templates, and significant improvements in motion correction and signal-to-noise ratio relative to previous studies (see Methods for further details).

We investigated the development of the DMN, DAN and ECN and of their relationship in neonates delivered and scanned at full-term (N = 282) relative to adults (N = 176). To understand the effect of neonate age on these networks, we also assessed preterm neonates. To deconfound the effect of the chronological age at the time of assessment from the effect of premature birth (Bhutta et al., 2002), we included two groups of preterm neonates: the first (N = 73) were scanned at TEA, and the second (N = 73) before TEA. We reasoned that any differences between neonates born and scanned at full-term and preterm neonates scanned at TEA would reflect effects of premature birth, while controlling for neonate age. Conversely, any differences between preterm neonates scanned at TEA and those scanned before TEA would reflect effects of neonate age.

## Methods

### Participants

#### Neonates

The neonate data were from the second (2019) dHCP public data release (http://www.developingconnectome.org/second-data-release/). All neonates were scanned at the Evelina Newborn Imaging Centre, Evelina London Children’s Hospital. Ethical approval was obtained from the UK’s National Research Ethics Committee and parental informed consent was obtained prior to imaging. *Full-term neonates*. We used 282/343 scans in the full-term neonates (gestational age (GA) at birth = 40.0 weeks ± 8.6 days; PMA at scan = 41.2 weeks ± 12.0 days; 160 males) after quality control procedures. *Preterm neonates*. We used 73 scans in both the preterm neonates scanned at TEA (GA at birth = 32.0 weeks ± 25.6 days; PMA at scan = 40.9 weeks ± 14.5 days; 41 males) and preterm neonates scanned before TEA (GA at birth = 32.5 weeks ± 13.4 days; PMA at scan = 34.6 weeks ± 13.4 days; 50 males). Of the 47 preterm neonates scanned both at and before TEA, 10 were discarded because of excessive movement of either one of the two scans, resulting in 37 paired scans (GA at birth = 31 weeks ± 6.8 days; PMA at first scan = 34 weeks ± 1.3 days; PMA at second scan: 40 weeks ± 6.9 days; 24 males). Further details in Figure 1, and SI file and Table S2.

**Figure 1.**
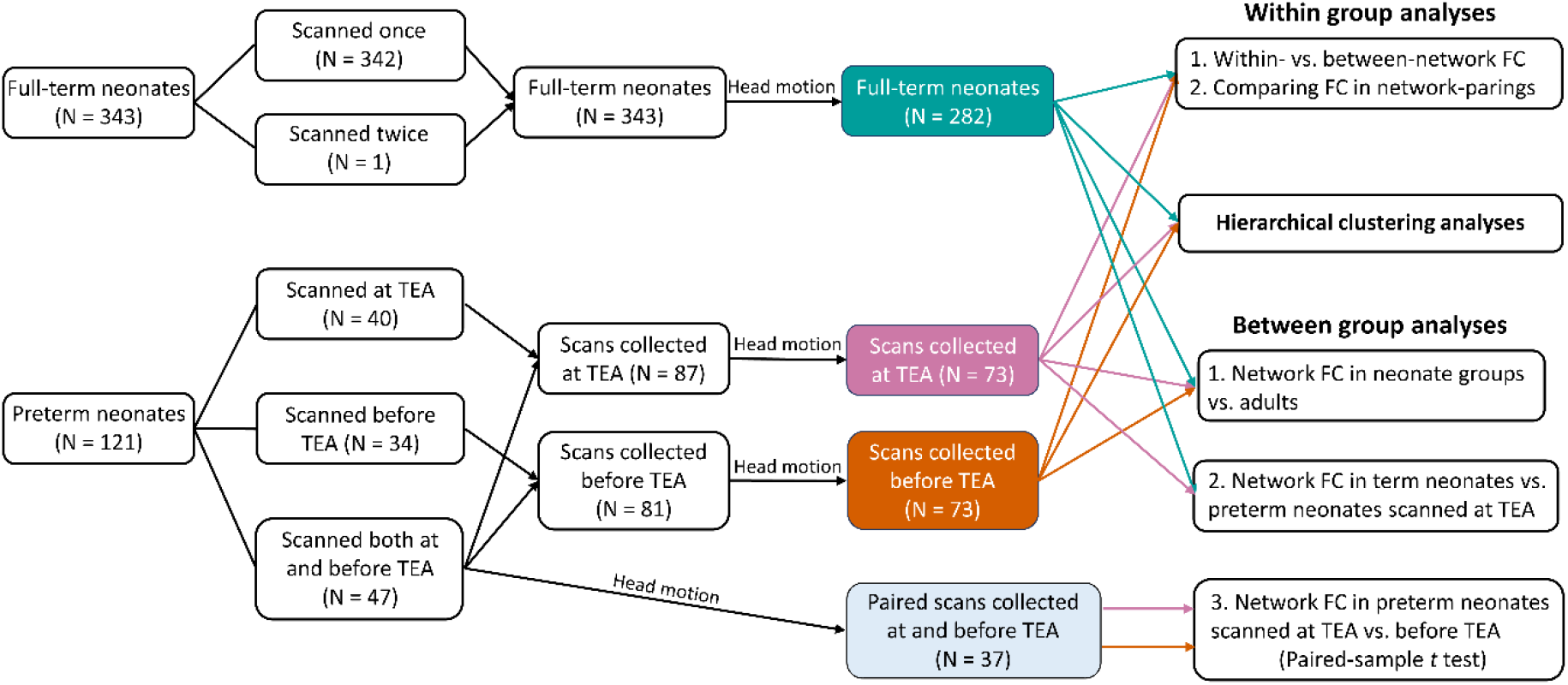
The number of scans included in data analyses. The frame in green/pink/brown indicate the scans collected from full-term neonates/preterm neonates at/before term-equivalent age that passed head motion criteria. The frame in light blue indicates the scans collected at and before term-equivalent age from the same preterm neonates. Abbreviations: TEA, term-equivalent age; vs., versus; FC, functional connectivity.

#### Adults

As a reference adult group, we used a subset (N = 176; 22 – 36 years; 77 males) of high quality data from the final release of the Washington University-Minnesota Consortium of Human Connectome Project (HCP) selected by Ito et al. (2020) (https://github.com/ito-takuya/corrQuench). For details of study procedures see Van Essen et al. (2013).

### Data acquisition and pre-processing

#### dHCP

Data were acquired on a 3T Philips Achieva with a dedicated neonatal imaging system including a neonatal 32-channel phased-array head coil. Fifteen minutes of high temporal and spatial resolution rs-fMRI data were acquired using a multislice gradient-echo echo planar imaging (EPI) sequence with multiband excitation (TE = 38 ms; TR = 392 ms; MB factor = 9x; 2.15 mm isotropic, 2300 volumes). In addition, single-band EPI reference (sbref) scans were also acquired with bandwidth-matched readout, along with additional spin echo EPI acquisitions with 4xAP and 4xPA phase-encoding directions. To correct susceptibility distortion in rs-fMRI data, field maps were also obtained from an interleaved (dual TE) spoiled gradient-echo sequence (TR = 10 ms; TE1 = 4.6 ms; TE2 = 6.9 ms; flip angle (FA) = 10°; 3 mm isotropic in-plane resolution, 6mm slice thickness). High-resolution T1- and T2-weighted anatomical imaging were also acquired in the same scan session, with a spatial resolution of 0.8 mm isotropic. For T1w image: TR = 4795 ms and the field of view (FOV) = 145 × 122 × 100 mm. For T2w image: TR = 12000 ms, TE = 156 ms and the FOV = 145 × 122 × 100 mm.

The dHCP rs-fMRI data were pre-processed by dHCP group using the project’s in-house pipeline optimized for neonatal imaging. See SI and Fitzgibbon et al. (2020) for full details. In order to reduce signal artefacts related to head motion, the cardiorespiratory fluctuations and multiband acquisition, the 24 extended rigid-body motion parameters together with single-subject ICA noise components were regressed out. To further reduce the effect of motion on functional connectivity measures, motion-outlier volumes were identified, and a scrubbing procedure was applied to retain a continuous sub-sample of the data (∼70%) with the lowest motion for each participant. The subjects who still had a high level of motion after scrubbing procedure were excluded from further analyses. We discarded the first 5 volumes to allow for adaptation to the environment and equilibrium of the MR signal at first. Then, motion outliers were identified from the remaining 2295 volumes. Volumes with DVARS (the root mean square intensity difference between successive volumes) higher than 1.5 interquartile range above the 75th centile, after motion and distortion correction, were considered motion outliers. Then, a continuous sub-sample of 1600 volumes with the minimum number of motion outliers was retained for each subject. Subjects with more than 160 motion-outlier volumes (10% of the cropped dataset) in the continuous subset were labelled ‘high level of motion’ and excluded entirely. Thus, 8 preterm neonates scanned before TEA, 14 preterm neonates scanned at TEA and 61 full-term neonates were excluded. In addition, we performed a temporal low-pass filter (0.08 Hz low-pass cutoff) on the pre-processed dHCP rs-fMRI to conduct functional connectivity (FC) analyses, as previous studies (Zuo et al., 2010; Salvador et al., 2008) found that oscillations were primarily detected within grey matter in 0.01 – 0.08 Hz. Figure S1a provides a schematic of the processing steps for dHCP fMRI data.

#### HCP

Data were acquired on a customized 3T Siemens “Connectome Skyra” with a 32-channel head coil. Resting state images were collected using gradient-echo EPI sequence: TR = 720 ms; TE = 33.1 ms; FA = 52°; FOV = 208 × 180 mm (RO × PE), slice thickness = 2 mm, 72 slices, 2.0 mm isotropic voxels, 1200 volumes per run. rs-fMRI data were pre-processed by HCP group. See SI and Van Essen et al. (2013) for full details.

### Data analyses

#### Network definition

We used a theory and meta-analyses driven node-based approach to network mapping. Nineteen regions of interest (8-mm radius spheres) for the three networks (Table S3), DMN, DAN and ECN, were created based on well-established landmark regions of interest (ROIs) defined in Raichle (2011). This method also helps to relate findings to our previous findings based on the same parcellation template (Naci et al., 2018; Haugg et al., 2018). See SI for details of alignment to neonate week-to-week structural templates and full details.

#### Functional connectivity (FC)

FC between ROIs was assessed by calculating the Pearson correlation of pre-processed time-courses, and z scored by using the Fisher-*z* transformation. At the individual level, within-network FC were obtained by averaging the FC between ROIs belonging to same network and between-network FC by averaging the FC between ROIs of each network to the others. For all between-group comparisons, the ROI-level FC within each subject was normalized to facilitate a focus on the FC-patterns between groups rather than potentially differing FC strength between the groups (Eyre et al., 2020; Smyser et al., 2016). ANOVAs and *t* tests were used to explore within and between group differences. Bonferroni correction for multiple comparison was applied to all statistical results.

#### Comparison of neonates and adults

General linear models (GLM) were used to test for group differences in FC within DMN, DAN or ECN while controlling for head motion, as we found neonates had significantly higher head motion than adults (SI, Figure S2, S3).

Hierarchical clustering analysis (Ripley et al., 2007; Rasmussen et al., 1992) and non-metric multidimensional scaling were used to capture network structure, and visualize the similarity of ROI responses in neonates and adults. See SI for further details.

#### Comparison of neonate groups

To investigate the effect of preterm birth, while controlling for age at scan, on network development, we compared the FC within each network and between each pair of networks, between preterm neonates scanned at TEA and full-term neonates. Head motion was not included as covariate, since we did not observe significant difference between the two groups (Figure S3). To investigate the effect of neonate age, while controlling for prematurity, the same preterm neonates scanned before and at TEA (N = 37) were used.

## Results

### The development of high-order networks in neonates

For full-term neonates, a 2 × 3 repeated measure ANOVA [type of FC (within-network, between-network) × network (DMN, DAN, ECN)] showed a significant main effect of type of FC (*F* (1, 281) = 766.14, *p* < 0.001), which was driven by higher overall connectivity for the within- relative to between-network connectivity (*t* (281) = 27.63, *p* < 0.001) (Figure 2a). We also found a main effect of network (*F* (2, 562) = 16.29, *p* < 0.001), which was driven by lower overall connectivity for the ECN relative to the DMN (*t* (281) = −3.06, *p* < 0.005) and DAN (*t* (281) = −5.84, *p* < 0.001). A significant interaction effect of type of FC by network (*F* (1.96, 550.18) = 81.44, *p* < 0.001) was driven by smaller difference between within- and between-network connectivity in the DMN relative to the DAN (*t* (281) = −4.41, *p* < 0.001), and in the ECN relative to the DMN (*t* (281) = −8.88, *p* < 0.001), and the DAN (*t* (281) = −11.86, *p* < 0.001). Paired-*t* tests showed significantly higher within- to between-network FC for each network (DMN: *t* (281) = 17.32, *p* < 0.001; DAN: *t* (281) = 21.05, *p* < 0.001; ECN: *t* (281) = 5.51, *p* < 0.001) (Figure 2a). This suggested that the coherence of nodes within each network was stronger than with the other networks’ nodes, in other words, that each of the three networks was differentiated as a cohesive unit, distinct from the other networks.

**Figure 2.**
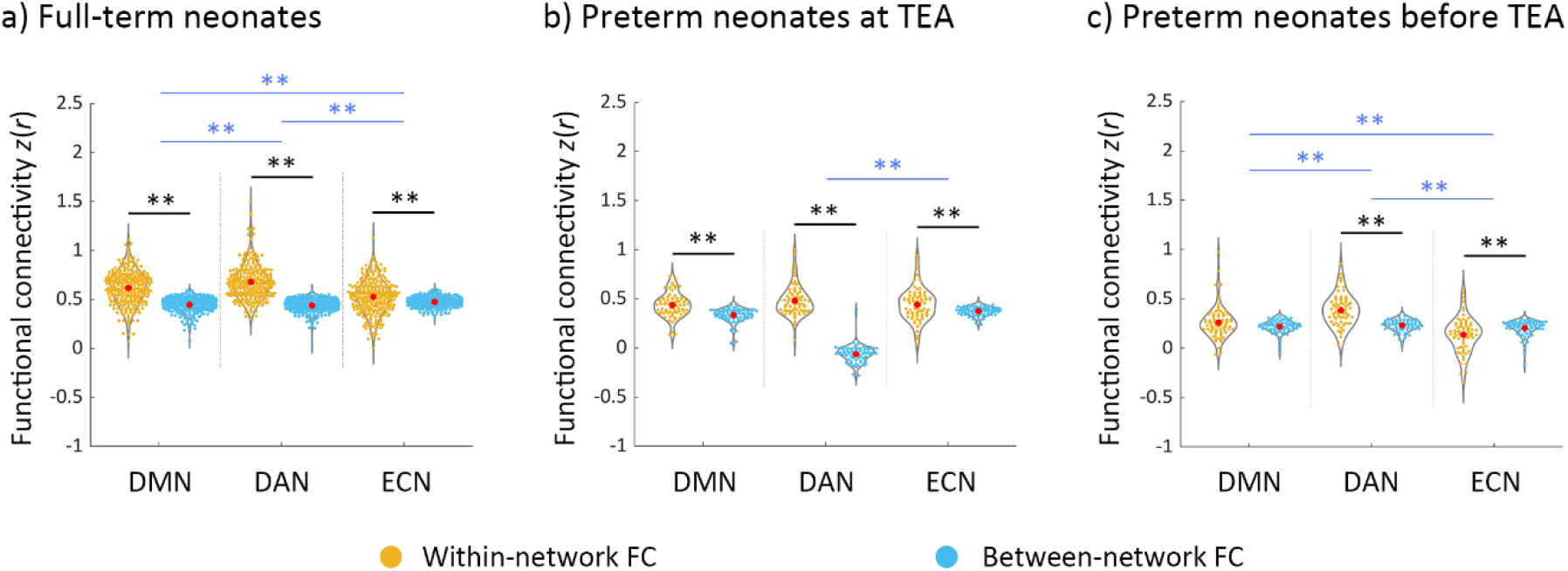
Within-network and between-network functional connectivity across DMN, DAN and ECN in the neonate groups. a) full-term neonates; b) preterm neonates scanned at term-equivalent age (TEA); c) preterm neonates scanned before TEA. The black lines/asterisks indicate significant difference between functional connectivity (FC) measures for each network, and the blue lines/asterisks indicate significant difference in FC measures of different networks. The FC values were Fisher-*z* transformed and inter-subject variability was removed for display purposes. Abbreviations: TEA, term-equivalent age; DMN, default mode network; DAN, dorsal attention network; ECN, executive control network; FC, functional connectivity; ** = *p* < 0.005.

For preterm neonates scanned at TEA, a similar 2 × 3 repeated measure ANOVA showed a significant main effect of FC type (*F* (1, 72) = 87.04, *p* < 0.001), which was driven by higher overall connectivity within- relative to between-network connectivity (*t* (72) = 9.35, *p* < 0.001) (Figure 2b). A significant interaction effect of type of FC by network (*F* (1.81, 130.23) = 6.03, *p* < 0.005), was driven by a smaller difference between within- and between-network connectivity in ECN relative to the DAN (*t* (72) = −2.96, *p* < 0.005). Paired-*t* tests showed significantly higher within- relative to between-network FC for each network (DMN: *t* (72) = 6.20, *p* < 0.001; DAN: *t* (72) = 8.05, *p* < 0.001; ECN: *t* (72) = 3.46, *p* < 0.001) (Figure 2b), suggesting that the DMN, DAN, and ECN were distinct from the one another in preterm neonates scanned at TEA.

For preterm neonates scanned before TEA, a similar 2 × 3 repeated measure ANOVA showed a significant main effect of type of FC (*F* (1, 72) = 16.80, *p* < 0.001), which was driven by higher overall connectivity for the within- relative to between-network connectivity (*t* (72) = 4.12, *p* < 0.001) (Figure 2c). A main effect of network (*F* (1.83,131.64) = 22.15, *p* < 0.001) was driven by lower overall connectivity for the DMN (*t* (72) = −3.98, *p* < 0.001) and ECN (*t* (72) = −6.45, *p* < 0.001) relative to the DAN (Figure 2c). A significant interaction effect of type of FC by network (*F* (2, 144) = 33.78, *p* < 0.001) was driven by smaller difference between within- and between-network connectivity in DMN relative to that in DAN (*t* (72) = −3.93, *p* < 0.001), and in ECN relative to that in DMN (*t* (72) = −4.32, *p* < 0.001) and DAN (*t* (72) = −8.21, *p* < 0.001). Paired-*t* tests showed significantly higher within- relative to between-network FC for DAN (*t* (72) = 7.73, *p* < 0.001), but significantly lower within-network FC compared to between-network FC for ECN (*t* (36) = −3.86, *p* < 0.001) (Figure 2c), suggesting that only the DAN was distinct from the other two networks in preterm neonates scanned before TEA.

In summary, these results suggested that the DMN, DAN and ECN were present in full-term and preterm neonates scanned at TEA, but only the DAN was formed as a distinct network in the preterm neonates scanned before TEA. Furthermore, the DAN was the most cohesive network in all three neonate groups.

### The development of the reciprocal relationship between the DMN and fronto-parietal networks in neonates

In full-term neonates, a one-way ANOVA with repeated measures for between-network FC (DMN–DAN, DMN–ECN, DAN–ECN) showed a significant main effect (*F* (1.98, 555.60) = 27.11, *p* < 0.001), which was driven by significantly lower FC in the DMN–DAN relative to DMN–ECN (*t* (281) = −7.79, *p* < 0.001) and DAN–ECN (*t* (281) = −4.64, *p* < 0.001) pairings. (Figure 3a). Similarly, to the adult data (see SI Results, Figure S4), the lower DMN–DAN FC, relative to the other pairings suggested that the reciprocal relationship between the DMN and DAN was present in full-term neonates. In preterm neonates scanned at TEA, a similar main effect (*F* (1.87, 134.45) = 11.93, *p* < 0.001), was driven by significantly lower FC in the DMN–DAN compared to DMN–ECN (*t* (72) = −4.78, *p* < 0.001) and DAN–ECN (*t* (72) = −4.20, *p* < 0.001) pairings (Figure 3b), and suggested that the reciprocal relationship between the two networks was present in preterm neonates scanned at TEA. In preterm neonates scanned before TEA, a significant main effect (*F* (2, 144) = 4.86, *p* = 0.009), was driven by lower FC in DMN–ECN relative to DMN–DAN (*t* (72) = −2.88, *p* = 0.005) and DAN–ECN (*t* (72) = −2.94, *p* < 0.005) pairings (Figure 3c). This is consistent with aforementioned results suggesting that the DMN and ECN are not yet developed as distinct networks in this group (Figure 2c). Furthermore, these results suggested that, by contrast to the full-term and preterm neonates scanned at TEA, the relationships between the three networks in preterm neonates scanned before TEA do not yet resemble the adult pattern (see SI, Figure S5). In summary, these results suggested that the reciprocal relationship between the DMN and DAN has started to develop in full-term neonates and preterm neonates scanned at TEA, but not in preterm neonates scanned before TEA.

**Figure 3.**
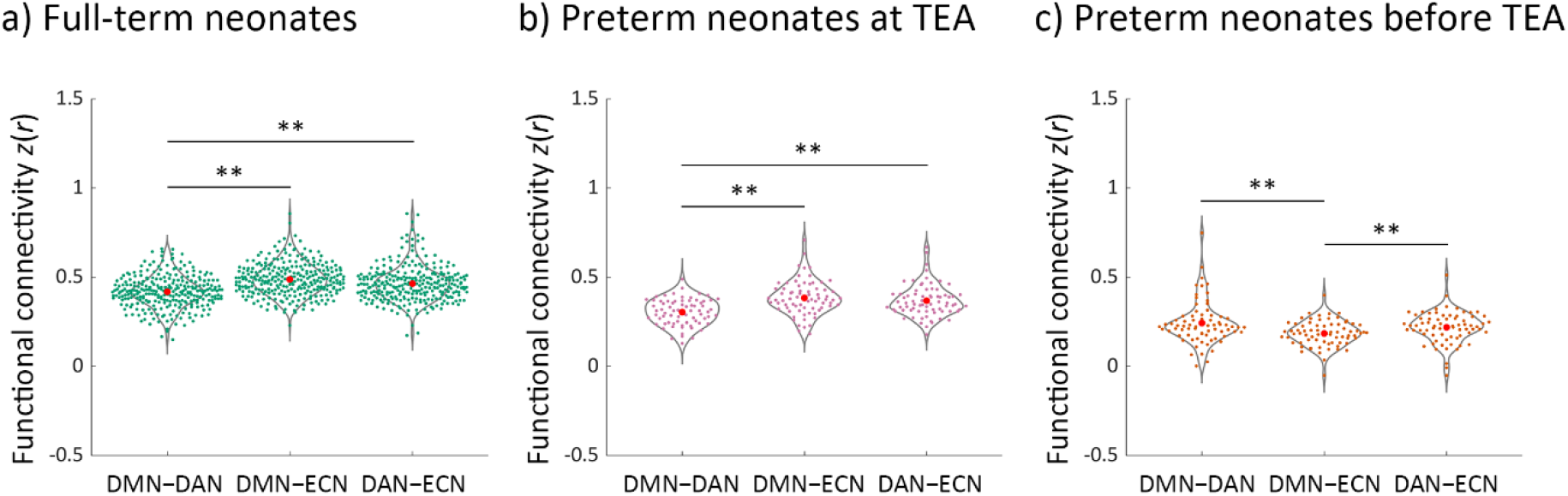
Between-network functional connectivity in neonate groups. a) full-term neonates; b) preterm neonates scanned at term-equivalent age (TEA); c) preterm neonates scanned before TEA. The FC values were Fisher-*z* transformed and inter-subject variability was removed for display purposes. Abbreviations: TEA, term-equivalent age; DMN–DAN, FC between the default mode network and dorsal attention network; DMN–ECN, FC between the default mode network and executive control network; DAN–ECN, FC between the dorsal attention network and executive control network; ** = *p* < 0.005.

### Comparison of neonate and adult networks

Visual inspection of the connectivity matrices (Figure 4) suggested that each neonate group had less cohesive networks (lower within relative to between-network connectivity) that the adult group. To investigate specifically how the three networks in neonates differed to those of adults, we compared the within- (Figures 5, 6) and between-network (Figure 7) connectivity in the neonate and adult groups.

**Figure 4.**
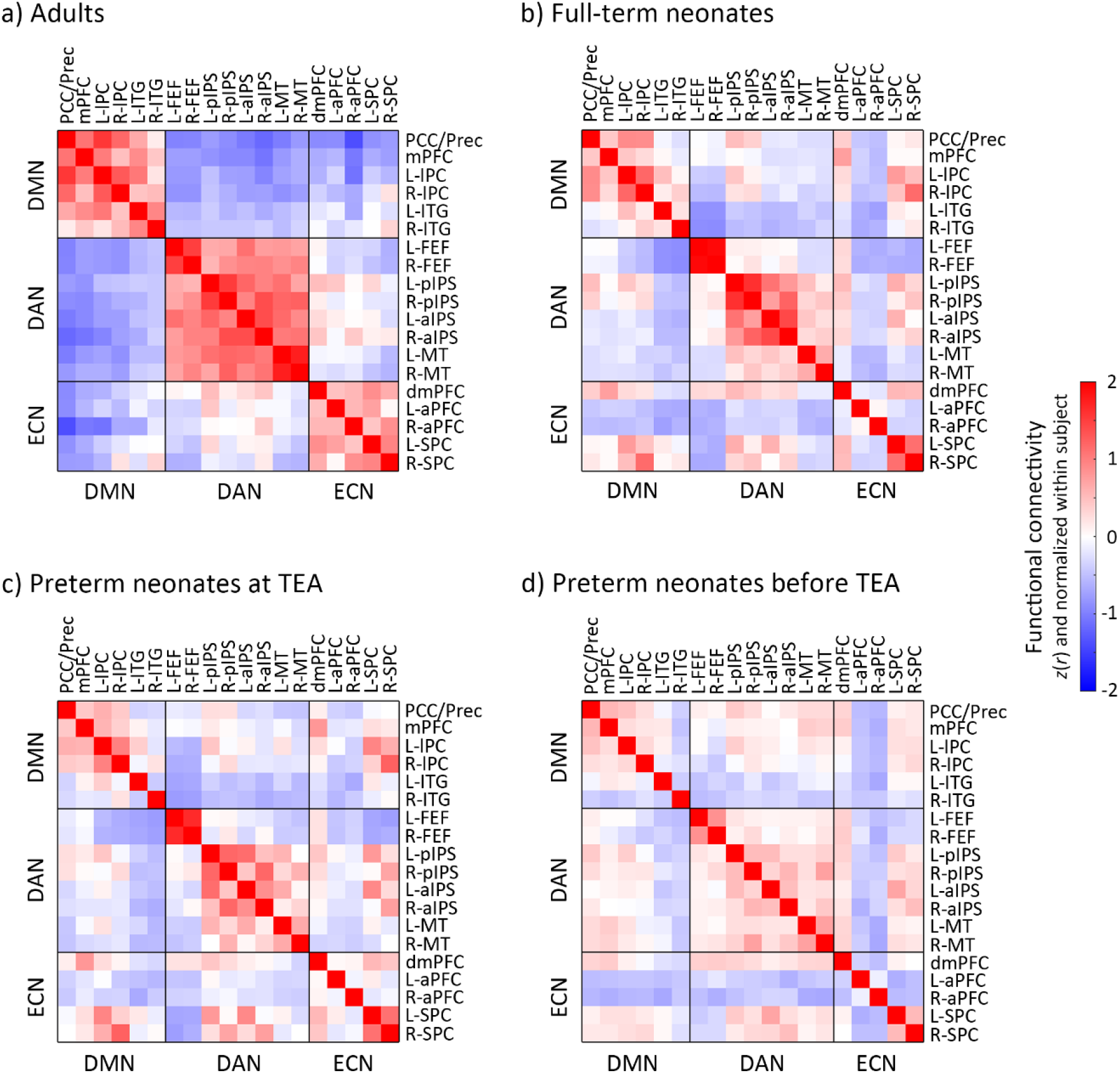
Functional connectivity (FC) in adults and neonate groups. a) adults, b) full-term neonates, c) preterm neonates scanned at term-equivalent age (TEA) and d) preterm neonates scanned before TEA. The FC value presents here was Fisher-*z* transformed and normalized within each subject before averaged within each group. Blue–red indicates low–high values. Abbreviations: TEA, term-equivalent age; DMN, default mode network; DAN, dorsal attention network; ECN, executive control network; R, right; L, left; PCC/Prec, posterior cingulate cortex/precuneus; mPFC, medial prefrontal cortex; lPC, lateral parietal cortex; ITG, inferior temporal gyrus; FEF, frontal eye field; pIPS, posterior intraparietal sulcus; aIPS, anterior intraparietal sulcus; MT, middle temporal area; dmPFC, dorsal medial prefrontal cortex; aPFC, anterior prefrontal cortex; SPC, superior parietal cortex.

**Figure 5.**
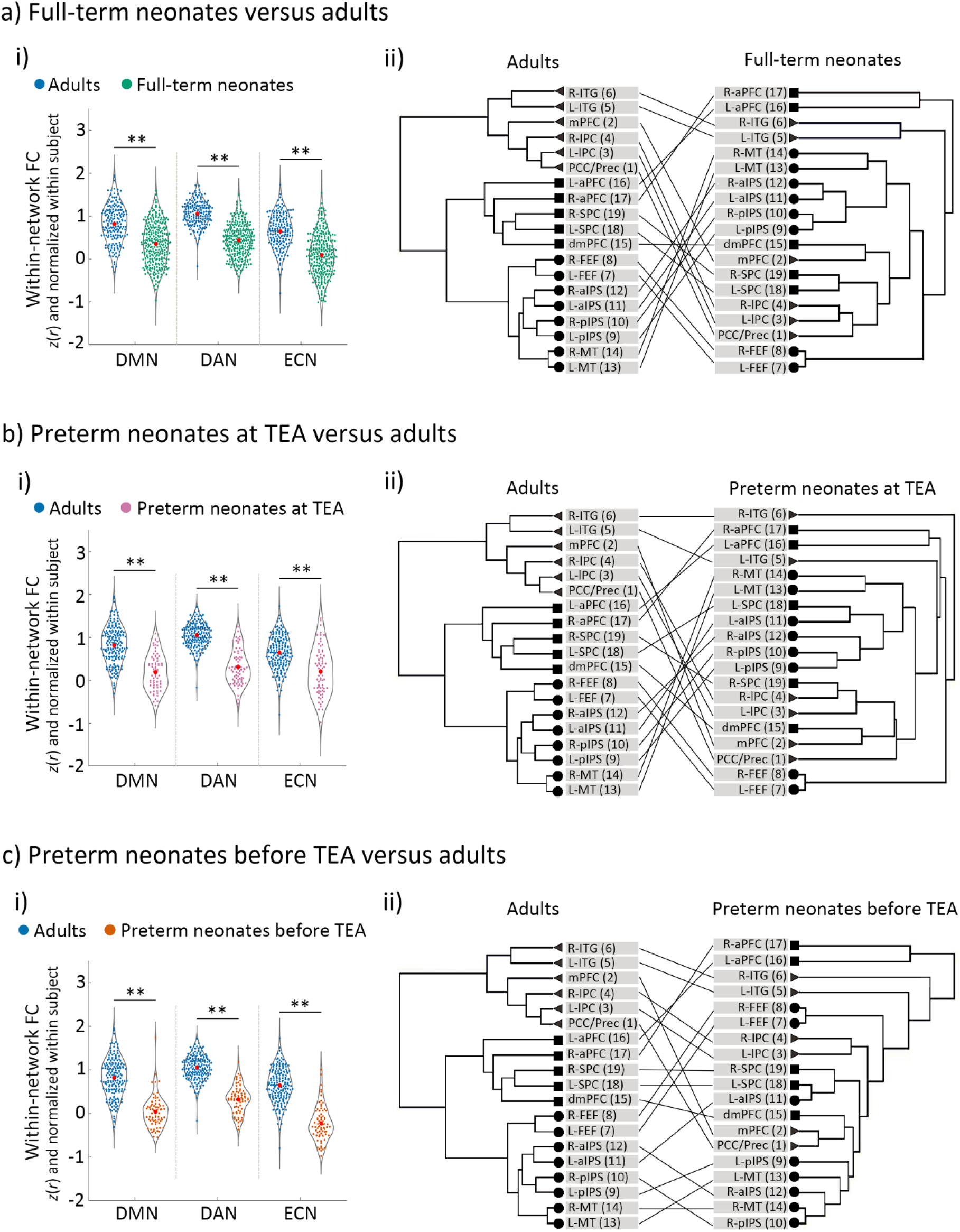
The development of the DMN, DAN and ECN in neonates relative to adults. a) full-term neonates, b) preterm neonates scanned at term-equivalent age (TEA) and c) preterm neonates scanned before TEA relative to the adults. Panels a-i), b-i) and c-i) depict the comparison of within-network functional connectivity (FC) between each neonate group and the adults. The FC values were Fisher-*z* transformed and normalized within each subject before averaging within each group. Panels a-ii), b-ii) and c-ii) depict the network structure of each neonate group relative to adults. Triangles/circles/squares represent the nodes of the DMN/DAN/ECN. Abbreviations: FC, Functional connectivity; DMN, default mode network; DAN, dorsal attention network; ECN, executive control network; R, right; L, left; PCC/Prec, posterior cingulate cortex/precuneus; mPFC, medial prefrontal cortex; lPC, lateral parietal cortex; ITG, inferior temporal gyrus; FEF, frontal eye field; pIPS, posterior intraparietal sulcus; aIPS, anterior intraparietal sulcus; MT, middle temporal area; dmPFC, dorsal medial prefrontal cortex; aPFC, anterior prefrontal cortex; SPC, superior parietal cortex; ** = *p* < 0.005.

**Figure 6.**
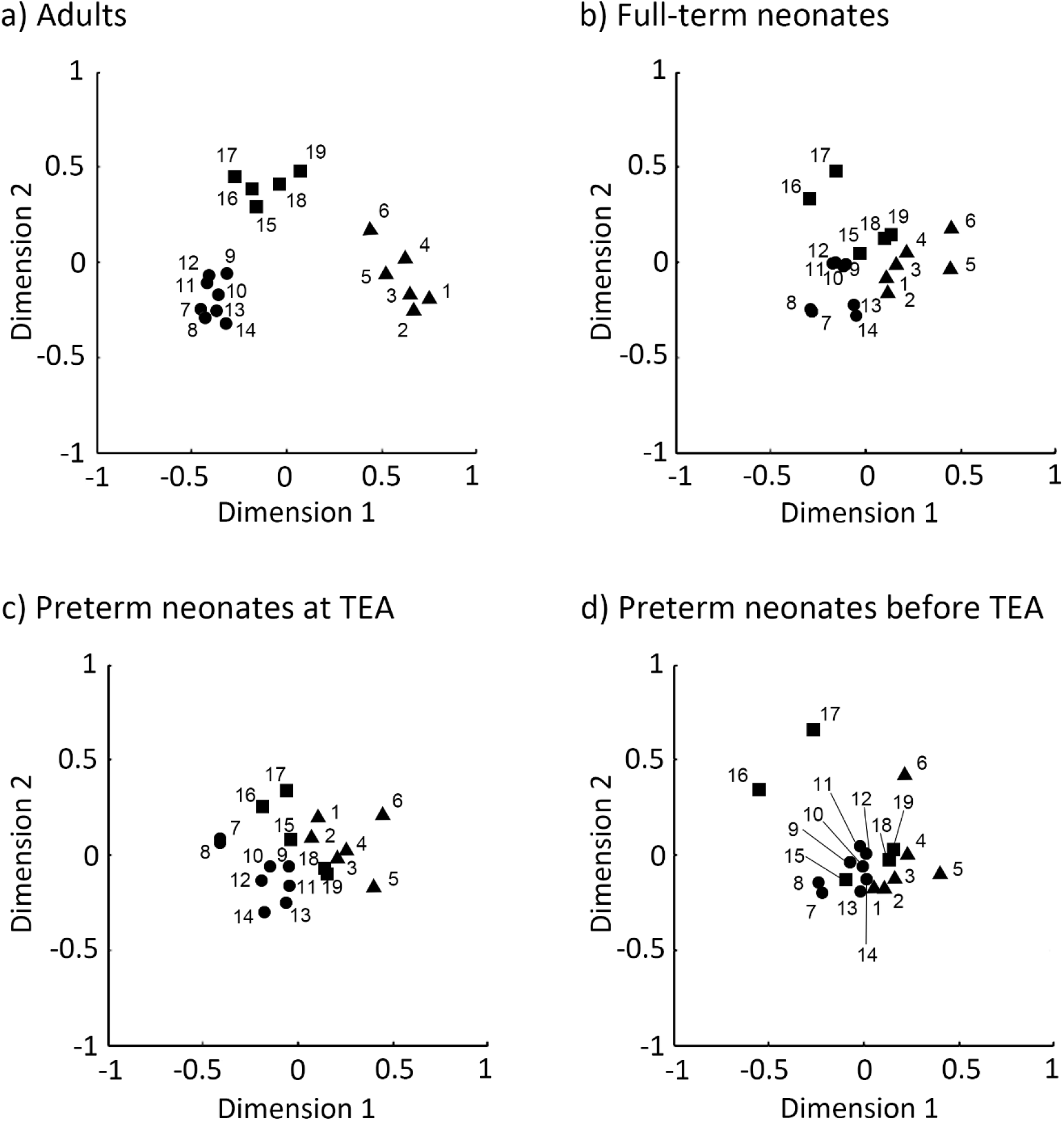
Multidimensional scaling (MDS) plots of regions in adults and neonates. a) adults, b) full-term neonates, c) preterm neonates scanned at term-equivalent age (TEA) and d) preterm neonates scanned before TEA. The 2-D plots were created using nonmetric MDS based on node’s similarity. Here triangles/circles/squares indicate nodes of the default mode/dorsal attention/executive control network. Abbreviations: 1, posterior cingulate cortex/precuneus; 2, medial prefrontal cortex; 3, left lateral parietal cortex; 4, right lateral parietal cortex; 5, left inferior temporal gyrus; 6, right inferior temporal gyrus; 7, left frontal eye field; 8, right frontal eye field; 9, left posterior intraparietal sulcus; 10, right posterior intraparietal sulcus; 11, left anterior intraparietal sulcus; 12, right anterior intraparietal sulcus; 13, left middle temporal area; 14, right middle temporal area; 15, dorsal medial prefrontal cortex; 16, left anterior prefrontal cortex; 17, right anterior prefrontal cortex; 18, left superior parietal cortex; 19, right superior parietal cortex; TEA, term-equivalent age.

**Figure 7.**
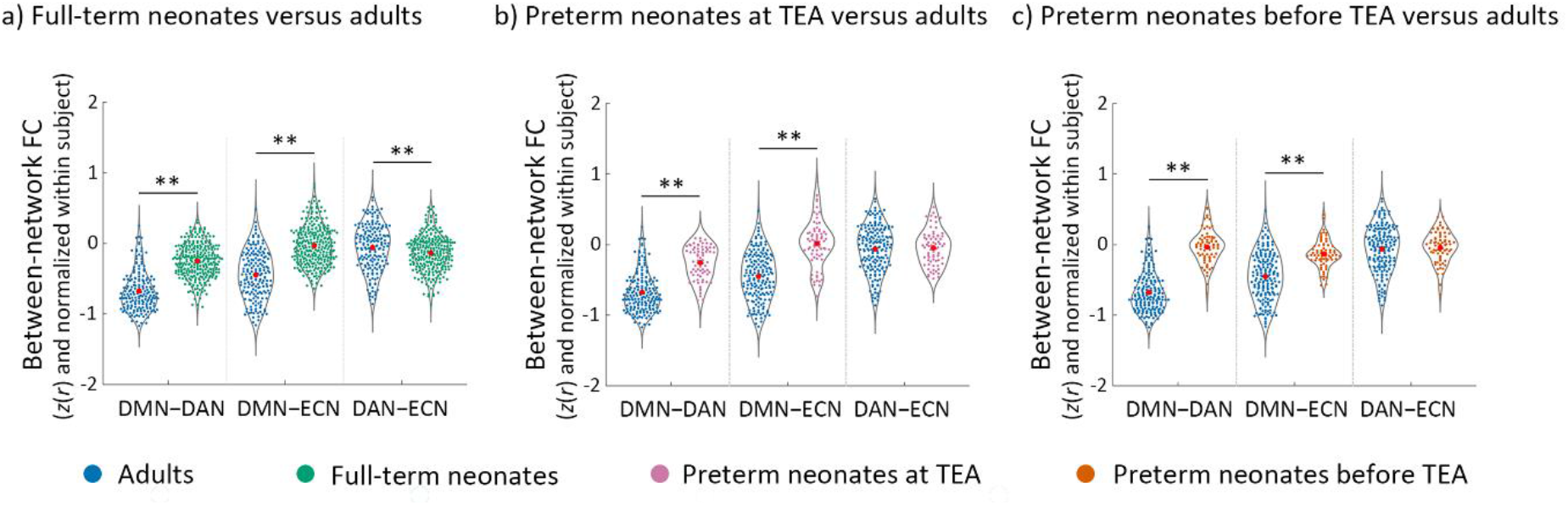
The development of the between-network functional connectivity (FC) in neonates relative to adults. a) full-term neonates, b) preterm neonates scanned at term-equivalent age (TEA) and c) preterm neonates scanned before TEA relative to the adults. The FC values were Fisher-*z* transformed and normalized within each subject before averaging within each group. Abbreviations: FC, functional connectivity; DMN–DAN, FC between DMN and DAN; DMN–ECN, FC between DMN and ECN; DAN–ECN, FC between DAN and ECN; ** = *p* < 0.005.

A GLM comparing adults and full-term neonates, including head motion as a covariate (SI, Figure S3), showed a significant main effects of group for all of the three networks (DMN: *F* (1, 455) = 75.62, *p* < 0.001); DAN: *F* (1, 455) = 333.33, *p* < 0.001); ECN: *F* (1, 455) = 135.88, *p* < 0.001); Figure 5a-i), which was driven by significantly higher within-network FC in the adults relative to full-term neonates. Hierarchical clustering analyses showed that, the adults’ network nodes grouped neatly into the a-priory postulated three distinct clusters (Raichle 2011), each comprising all the ROIs belonging to that network (Table S3). By contrast, in the full-term neonates, the ROIs clustered into groups that were inter-mixed between the three networks (Figure 5a-ii). Similarly, for the preterm neonates scanned at TEA, we found significant main effects of group for all of the three networks (DMN: *F* (1, 246) = 90.16, *p* < 0.001); DAN: *F* (1, 246) = 252.32, *p* < 0.001); ECN: *F* (1, 246) = 42.40, *p* < 0.001); Figure 5b-i), again driven by significantly higher within-network FC in the adult group. Unlike the adult group, preterm ROIs clustered into groups inter-mixed between the three networks (Figure 5b-ii). Consistent with the other two groups’ results, for the preterm neonates scanned before TEA, we found significant main effects of group for all of the three networks (DMN: *F* (1, 246) = 145.15, *p* < 0.001); DAN: *F* (1, 246) = 276.15, *p* < 0.001); ECN: *F* (1, 246) = 218.91, *p* < 0.001); (Figure 5c-i)), which were driven by significantly higher within-network FC in adults, and ROI clusters that did not adhere to network identity (Figure 5c-ii).

Multidimensional scaling analyses further confirmed these results, by showing that the adult ROIs formed three distinct cluster conforming to network identify (Figure 6a), distanced from one another in representational space. By contrast, each network’s ROIs in the neonate groups formed less distinct clusters, i.e., clusters were more closely grouped together, and their separability as distinct clusters was reduced with neonate age. The preterm neonates scanned before TEA showed the most intermingling of ROIs across the three networks in this two-dimensional manifold (Figure 6d).

In summary, these results suggested that although the three networks were present in full-term and preterm neonates scanned at TEA, and the DAN in preterm neonates scanned before TEA, their coherence was lower and structure less well-organised than the canonical adult networks. This is consistent with previous studies showing that brain networks continue to develop from birth onwards (Turk et al., 2019; Keunen et al., 2017; Cusack et al., 2016; Doria et al., 2010; Limperopoulos et al., 2005). As expected, the reciprocal relationship between the DMN and fronto-parietal networks was also weaker in neonates relative to adults (Figure 7) (see SI Results for full details).

### The effect of prematurity on network development and their relationship

Comparison of network FC in full-term relative to preterm neonates scanned at TEA showed significantly lower FC within the DMN (*t* (353) = −2.64, *p* = 0.009) and the DAN (*t* (353) = −2.63, *p* = 0.009) in the preterm group (Figure 8a). In addition, we observed higher FC between the DAN and ECN (*t* (353) = 2.83, *p* = 0.005) in preterm neonates. These results suggested that, by term-equivalent age, preterm birth is associated with lower DMN and DAN network coherence, and lower differentiation of the DAN and ECN from one another, relative to full-term birth.

**Figure 8.**
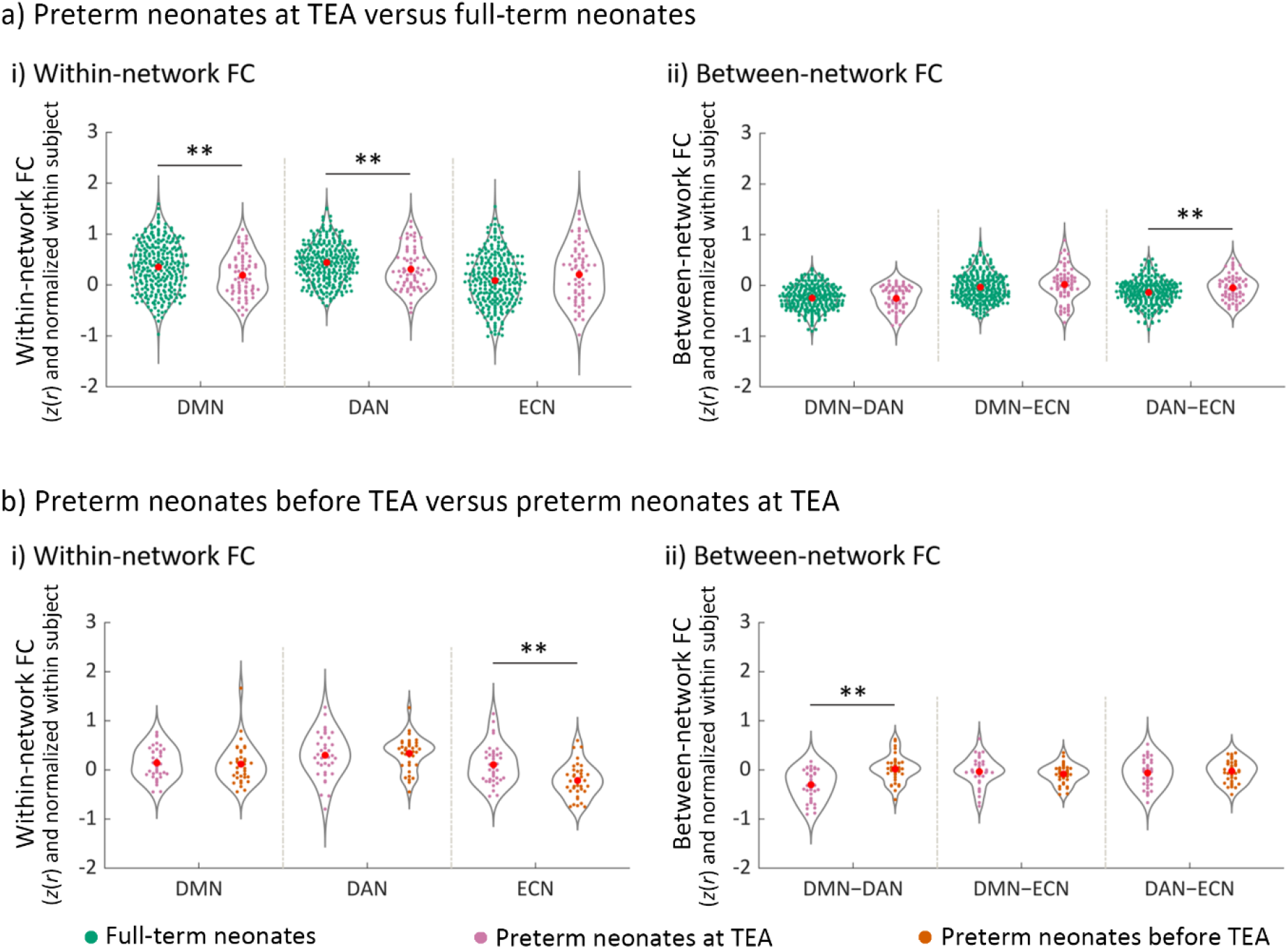
The effect of premature birth and early neonate age on the development of network functional connectivity (FC). a) The effect of premature birth on a-i) within-network FC and a-ii) between-network FC. The FC values represented in a-i) and a-ii) were Fisher-*z* transformed and normalized within each subject before averaging within each group. b) The effect of early neonate age on b-i) within-network FC and b-ii) between-network FC. The FC values represented in b-i) and b-ii) were Fisher-z transformed, normalized within each subject, and inter-subject variability was removed for display purposes. Abbreviations: DMN, default mode network; DAN, dorsal attention network; ECN, executive control network; DMN–DAN, FC between DMN and DAN; DMN–ECN, FC between DMN and ECN; DAN–ECN, FC between DAN and ECN; ** = *p* < 0.005.

### The effect of premature neonate age on network development and their relationship

We found significantly lower FC within the ECN (*t* (36) = −4.28, *p* < 0.001) and higher DMN–DAN FC (*t* (36) = 4.39, *p* < 0.001), suggesting lower ECN network coherence and lower functional differentiation between the DMN and DAN, for preterm neonates scanned before TEA relative to the same group scanned at TEA (Figure 8b). Thus, these results suggested that neonate age, and particularly, development up to term-equivalent age, is a significant factor for the maturation of the ECN and of the development of the reciprocal relationship between the DMN and the DAN.

## Discussion

In this study, we asked whether infants have the capacity for conscious experiences. The lack of language and wilful motoric output present major barriers to consciousness science in neonates. To sidestep these limitations, we examined whether or not the brain circuitry of conscious awareness is developed at birth. In particular, we focused on the impact of prematurity and early infant age on the development the default mode and fronto-parietal networks, and of their reciprocal relationship. The critical novel contribution of this study is showing that the DAN, ECN and DMN are already present in neonates by full-term or term-equivalent age, and furthermore, that the reciprocal relationship between the DMN and the DAN, is already instated by this age. By contrast, this relationship is not present in preterm neonates before term-equivalent age. Our results in full-term neonates are consistent with a number of previous studies that observed high-order networks in this group (Rajasilta et al., 2020; Linke et al., 2018; He et al., 2016, 2015; Doria et al., 2010)

### Effect of preterm birth

We found that the three networks were present in preterm neonates at TEA. Furthermore, for the first time, we show that the reciprocal relationship between the DAN and DMN was instated in preterm neonates at TEA. Previous rs-fMRI neonate studies that have investigated the effect of preterm birth have mainly focused on disrupted within-network coherence (Eyre et al., 2020; Smyser et al., 2016) or topological organization (Cao et al., 2017; van den Heuvel et al., 2015), and knowledge of the impact of preterm birth on the relationship between high-order brain networks remains scarce. It is, therefore, striking to see that this key functional relationship develops in healthy-born premature neonates by term-equivalent age, according to a pre-programmed developmental trajectory despite of prematurity.

However, premature birth had a negative impact on network development. Preterm neonates at TEA had significantly lower within-network connectivity in the DMN and DAN, suggesting that these networks were less developed relative to full-term neonates despite the matched age. Furthermore, the DAN–ECN connectivity was higher in preterm neonates at TEA, likely due to weaker within-network connectivity in the DAN. Our results are consistent with Smyser et al. (2016) and Eyre et al. (2020) findings of lower within-network connectivity in preterm relative full-term neonates. Bouyssi-Kobar et al., (2019) showed decreased density of connections in parietal-temporal and frontal areas in preterm neonates scanned at TEA relative to full-term neonates, using graph theoretical modelling. Previous structural MRI studies (Bouyssi-Kobar et al., 2018; Pandit et al., 2014) also found widespread deficiencies in grey and white matter, including in the DMN regions, in preterms scanned at TEA relative to full-term neonates (Bouyssi-Kobar et al., 2018). Consistent with a growing literature, our results shed light on disrupted brain mechanisms that may underlie the significant risks for neurodevelopmental and psychiatric problems in later life (Bhutta et al., 2002; Marlow et al., 2005; Saigal and Doyle, 2008; Nosarti et al., 2012), that are associated with preterm birth.

### Impact of premature neonate age on network development

In contrast to preterm neonates assessed at TEA, neonates who were assessed before TEA showed dramatic underdevelopment of networks and their relationships. Only the DAN, but not the ECN and DMN, was present as a discrete network. The ECN showed significant development in the early weeks post premature birth; within-network connectivity became significantly stronger as premature neonates reached TEA. Strengthening of within-network connectivity in fronto-parietal regions has been reported in previous studies that assessed preterm neonates longitudinally (He et al., 2016). These findings suggest that, relative to other high-order networks, e.g., the DMN, the ECN aspect of the fronto-parietal networks is the least developed in preterms and most liable to undergo developmental change in the early weeks post birth. Furthermore, the reciprocal relationship between the fronto-parietal and DMN networks also strengthened in the weeks up to term-equivalent age. It was not formed before TEA, but appeared once preterms reached TEA. These results present a novel finding for premature neonates. They resonate with fetal studies (Thomason et al. 2015; 2014) showing that coactivation from the posterior cingulate cortex (node of the DMN) to areas of dorsal attention/executive networks became more negatively coupled with increasing fetal age.

### Comparison of neonate and adult networks

It is important to note that although the DAN, ECN and DMN were already present at full-term and term-equivalent age, they were significantly different from the adult networks. The neonate networks had significantly lower within-network connectivity and were atypical in their nodal structure, suggesting less within-network cohesiveness compared to the adults’. Similarly, the reciprocal relationship between the DMN and DAN was less developed in neonates relative to adults. These findings are consistent with previous studies (Sherman et al., 2014; Gao et al., 2009; Fair et al., 2007) showing that the functional organization of the brain rapidly develops from birth onwards. For example, Gao et al. (2009) reported that the DMN becomes adultlike by 2 years of age. Studies have also reported significant differences in segregation and integration, indicators of functional network maturation, in the DMN and ECN between childhood and adulthood (Sherman et al., 2014; Fair et al., 2007; 2008)

Of the three high-order networks, the ECN was the least developed in premature neonates, whereas the DAN was the most well-formed across all neonate groups. These results suggest different ontogenesis trajectories for the two fronto-parietal networks, likely explained by their differential functions. The DAN serves to orient and modulate attention to the saliency of incoming sensory inputs (Corbetta and Shulman 2002), a capacity that probably emerges early in fetal development, as foetuses start to perceive sounds inside the womb. By contrast, behavioural response planning and monitoring, subserved by the ECN (Kroger et al., 2002), is a mental faculty that relies heavily on the increasingly complex interactions with their environment that neonates engage in as they mature from the first weeks and months from birth.

### Methodological considerations

Although some previous studies have observed high-order networks in neonates (Rajasilta et al., 2020; Linke et al., 2018; He et al., 2015, 2016; Doria et al., 2010) others have not (Gao et al., 2015a, 2015b; Fransson et al., 2007; Eyre et al, 2020). This inconsistency is likely due to methodological differences in the type of data acquired and the analyses method. By contrast to some previous studies (Gao et al., 2015a, 2015b; Fransson et al., 2007), here we were enabled by the high quality dHCP dataset to employ a uniquely large sample of high temporal and spatial resolution infant rs-fMRI data, and accurate week-by-week structural templates of the developing infant brain, which ensured higher sensitivity to detect brain networks in early infancy. Second, we used a theoretically motivated region-of-interest analysis rather than a data-driven independent component analysis (ICA) for network definition. While ICA is a convenient data-driven tool to extract networks, it requires a critical free parameter that determines how many will be extracted, and hence the degree to which brain networks are likely to be detected as whole versus broken into parts. For example, a recent paper by Eyre et al. (2020), that used the dHCP dataset, did not find a single network corresponding to the whole DMN, but it is possible that this result would have changed with a different parameter choice. Given these methodological strengths and clear results, we believe this study helps to resolve previous inconsistent findings on the development of high-order brain network in neonates.

### Implications for understanding conscious awareness in neonates

What are the implications of our findings for understanding the conscious experiences of neonates? For the first time, here we show that by full-term birth or term-equivalent age, neonates possess key features of the brain infrastructure that enables the integration of information across diverse sensory and higher-order functional modules, which gives rise to conscious access. We also show that this system is yet to undergo substantial change before it resembles that of the adults. Therefore, while these findings suggest that the capacity for conscious experiences is present at birth, coupled with previous evidence, they suggest that such experiences may be limited. The frontal cortex, a nexus of the fronto-parietal and DMN networks, undergoes dramatic maturation and reorganization by the end of the first year of life, resulting in big improvements in several cognitive abilities, and in sophistication of related mental content at that age (Diamond and Goldman-Rakic, 1989). Consistent with neuroanatomical data, Kouider et al. (2013) found an electrophysiological signature of perceptual consciousness which was present albeit weak and delayed in 5-month-olds, became stronger and faster in 12–15-month-old infants. Kovács et al. (2010) found that infants’ eye movements demonstrated a capacity to monitor other people’s beliefs at 7 months, and to make predictions about visual scenes at 12 months. Critically, at birth, neonates are largely bereft of the wealth of prior experiences that inform episodic memory and semantic knowledge, which allow for the construction of longitudinal awareness (Seely and Sturm, 2006; Damasio, 2000), and the creation of an internal world that projects beyond the present and is extended across time. Nevertheless, even from the first days of life, neonates encode, store, and retrieve information about events in their world (Ellis et al., 2020; Howe et al., 2004; 2003). They start to integrate sensory, kinesthetic and proprioceptive stimulus response contingencies, in order to understand the actions of others and generate models for producing similar actions. Our results suggest that from birth neonates possess the capacity to integrate sensory and incipient cognitive experiences into coherent conscious experiences about their core self and the developing relationship to their environment.

## Acknowledgements

H.H. was funded by the China Scholarship Council – Trinity College Dublin Joint Scholarship Programme. R.C. is supported by an ERC Advanced Grant 787981 (FOUNDCOG). L.N. was funded by an L’Oreal for Women In Science International Rising Talent Award, and the Welcome Trust Institutional Strategic Support Fund. Neonate data were obtained from the Developing Human Connectome Project website (http://www.developingconnectome.org/second-data-release/) and adult data were obtained through the Washington University-Minnesota Consortium of Human Connectome Project website (http://www.humanconnectomeproject.org).

## Author contributions

Study Conceptualization, L.N., H.H.; Methodology, L.N., H.H., R.C.; Formal Analyses, H.H.; Writing, H.H., L.N.; Revision: R.C., L.N.; Funding and resource acquisition, L.N. Project administration, L.N.; Supervision, L.N.

