## Supplementary Information for "Typical and disrupted brain circuitry for conscious awareness in full-term and preterm infants"

**Title**

### Supplementary Methods

#### Participants

*Full-term neonates*: Data from 342 full-term neonates were available. Only one subject was scanned twice, at 37 weeks postmenstrual age (PMA) and after 37 weeks PMA, but we only used the later one for the full-term group. 61 scans were discarded because of excessive movement (see Data Analyses section). Thus, the full-term neonate group had 282 participants (gestational age (GA) at birth = 40.0 weeks ± 8.6 days; PMA at scan = 41.2 weeks ± 12.0 days; 160 males).

*Preterm neonates*. 121 preterm neonates had fMRI data in second dHCP public data release. 74 of them were scanned once: 40 participants scanned at TEA and 34 participants scanned before TEA. 47 of the 121 preterm neonates were scanned twice, at and before TEA. Thus, for preterm neonates, we have 87 obtained at TEA and 81 before TEA. 14/87 scans collected at TEA and 8/81 scans collected before TEA were discarded because of excessive movement (see Data Analyses section).

*Adults.* All participants were right-handed, native English speakers, and no history of neurological disorders. All participants were tested at Washington University in accordance with the protocol approved by the Washington University institutional review board. Informed consent was obtained for each participant prior to the experiment. Specific details and procedures of subject recruitment can be found in Van Essen et al. (2013). The subset used in current study passed stringent quality control measures relative to the larger HCP (Ito et al. 2020). Detailed information about exclusion criteria can be found in Ito et al. (2020) and the full list of the 176 participants used in this study is available here: https://github.com/ito-takuya/corrQuench.

#### Data Acquisition

*dHCP data acquisition.* Full details of dHCP data acquisition can be found at Fitzgibbon et al. (2020) Prior to scanning, neonates were fed, swaddled, and comfortably positioned in a vacuum jacket to promote natural sleep. Fifteen minutes of high temporal and spatial resolution resting state fMRI (rs-fMRI) data were acquired using a multislice gradient-echo echo planar imaging (EPI) sequence with multiband excitation (TE = 38 ms; TR = 392 ms; MB factor = 9x; 2.15 mm isotropic, 2300 volumes).

*HCP.* Full details of HCP data acquisition can be found at Van Essen et al. (2013). Rs-fMRI data were collected in four runs of 14.4 minutes each, two runs in one session and two in another session. The two sessions were conducted in two days separately and we only used the data collected in the first session. Within each session, oblique axial acquisitions alternated between phase encoding in a right-to-left (RL) direction in one run and phase encoding in a left-to-right (LR) direction in the other run. Resting state images were collected using gradient-echo EPI sequence: TR = 720 ms; TE = 33.1 ms; FA = 52°; FOV = 208 × 180 mm (RO × PE), slice thickness = 2 mm, 72 slices, 2.0 mm isotropic voxels, 1200 volumes per run.

**Data pre-processing**

*dHCP.* The dHCP rs-fMRI data were pre-processed by dHCP group using the project's in-house pipeline optimized for neonatal imaging and specifically developed for this dataset, detailed in Fitzgibbon et al. (2020). This pipeline includes: 1) Motion and distortion correction: corrects for intra-volume movement artefacts and for artefacts associated with susceptibility-induced off-resonance field changes (susceptibility-by-movement artefacts), and estimates motion nuisance regressors; 2) Registration: aligns all functional images with the native T2 space and the neonatal template space, which refers to the week-specific 40-week template from the dHCP volumetric atlas (Schuh et al., 2018); 3) Temporal high-pass filter: 150s high-pass cut-off; 4) Denoising: Estimates artefact nuisance regressors and regresses all nuisance regressors from the functional data obtained from the first step.

*HCP.* The rs-fMRI data of HCP were pre-processed by HCP group using following pipeline: 1) Distortion correction: correction of gradient-nonlinearity-induced distortion and phase-encoding-direction-induced distortion; 2) Motion correction: realigns the timeseries to correct for subject motion by using a 6 DOF FLIRT (Oxford Centre for Functional MRI of the Brain Fs Linear Registration Tool) registration of each frame to the single-band reference image; 3) Aligns the original EPI data (rs-fMRI data) to Montreal Neurological Institute (MNI) template space: EPI to T1w from FLIRT BBR, fine tuning of EPI to T1w with BBR-register, nonlinear T1w to MNI template; 4) Intensity normalization to mean of 10000 and bias field removal; 5) Temporal high-pass filter: 150s high-pass cut-off; 6) Denoising: removes artefactual or "bad" components using ICA-FIX to automatically. Detailed pre-processing procedure can be found in Glasser et al. (2013). Additionally, we performed a temporal low-pass filter (0.08 Hz low-pass cutoff) on the denoised rs-fMRI data and removed the first five volumes. Figure S1 provides a schematic of the processing steps for HCP fMRI data. As the selection of the subset of HCP had controlled head motion (i.e., exclusion of participants that had any fMRI run in which more than 50% of TRs had greater than 0.25mm framewise displacement), and adults generally have smaller maximal head motion than neonates (Cusack et al., 2017), we did not apply the same scrubbing method used in the dHCP dataset to adult data. To assess the effect of this, we compared the mean framewise displacement (FD) value (Power et al., 2014; Power et al., 2012) in adults and neonates before/after the scrubbing procedure. The FD value indexes the movement of the head from one volume to the next and is defined as the sum of the absolute values of the differential realignment estimates (the six realignment parameters). It has been widely used to index head movement and exclude subjects of high motion (Sripada et al., 2019; Gu et al., 2018; Hoptman et al., 2012). Independent-samples *t*-tests showed that adults had significantly lower head motion compared to neonates before ($t$ (491.45) = -12.49, $p < 0.001$) and even after scrubbing ($t$ (580.56) = -9.92, $p < 0.001$; Figure S2).

*Figure S1 about here please*

**Network definition**

We first aligned these ROIs with 40-week dHCP T1w template (Cabral et al., 2020). This involved: 1) trimming the dura from the 40-week dHCP T1w template, and the cerebellum from both the 40-week dHCP T1w template and MNI T1w template; 2) aligning the 40-week dHCP T1w template to MNI T1w template using non-linear registration (ANTs SyN); 3) applying the warp file generated in the last step to the ROIs in MNI space with 40-week dHCP T1w template as reference. In the next step, we needed to align these ROIs in 40-week dHCP T1w template space with neonate native space. We inverted the func-to-template warp provided by dHCP group and applied this inverted warp to ROIs in the 40-week dHCP T1w template space. Thus, we obtained ROIs in each neonate native functional space. For adults, as the denoised HCP data had been aligned to MNI space, we used these ROIs in MNI space directly.

**Data analyses**

***Hierarchical clustering analyses***. We captured the structure of the three networks in different groups with hierarchical clustering analysis (Ripley et al., 2007; Rasmussen et al., 1992), which has proven informative in prior infant studies (Cusack et al., 2018). This hierarchical clustering algorithm builds up an entire cluster tree in which neighbouring regions are joined if their similarity is maximal among all pairs of neighbouring regions. Here, we used the time-course extracted from the 19 ROIs as input to access the hierarchical relationship among the ROIs. For the neonate data, we first calculated initial pairwise distance between ROIs using one minus the linear correlation between the scrubbed time-courses extracted from the 19 ROIs at the individual level. For adults, the initial pairwise distance between ROIs were calculated using one minus the linear correlation between the time-courses of 1195 time points extracted from the 19 ROIs at the individual level. Then, we averaged the pairwise distances between ROIs within each group to get the group-level pairwise distances, which were submitted to hierarchical clustering analysis to create a hierarchical cluster tree of the 19 ROIs for each group respectively. The cophenetic correlation coefficient was used to create a dendrogram for each group. The length of each C link in the dendrogram represents the distance between regions/clusters.

***Multidimensional scaling analysis.*** Non-metric multidimensional scaling (MDS) was also used to facilitate visualizing the similarity of ROIs functional response for adults and neonate groups. The non-metric MDS (Kruskal, 1964) performs non-metric multidimensional scaling on the dissimilarity matrix of item−item (i.e., ROI−ROI dissimilarity matrix) to compute a configuration. Then, the Euclidean distances between items (i.e., ROIs) in the configuration were obtained. The difference between the monotonic transformed dissimilarities in the item−item (i.e., ROI−ROI) matrix and the Euclidean distances between items (i.e., ROIs) in this configuration were minimized and items (ROIs) were represented in a low-dimensional space (i.e., a 2-D space). The ROI−ROI dissimilarity matrix (one minus the linear correlation between the time-courses) for each group from the hierarchical clustering analysis was submitted to non-metric MDS analysis implemented in MATLAB.

### Supplementary Results

***Comparison of head motion in neonates and adults.*** Independent-sample *t*-tests were applied to compare the head motion in the adults and neonates before/after scrubbing procedure. We found that adults had significantly lower head motion compared to neonates before ($t$ (491.45) = -12.49, $p < 0.001$) and after ($t$ (580.56) = -9.92, $p < 0.001$) scrubbing procedure (Figure S2). A paired-*t* test was applied to detect the difference in head motion in neonates before and after scrubbing procedure. Results showed that neonates had significantly lower head motion after scrubbing relative to before scrubbing ($t$ (427) = -9.69, $p < 0.001$) (Figure S2). Bonferroni correction for multiple comparison was applied to statistical results.

*Figure S2 about here please*

***Comparison of head motion in neonate groups after scrubbing and adults.*** We conducted a one-way ANOVA to compare the difference in head motion between the adults and neonate groups after scrubbing procedure and found a significant main effect of group ($F$ (3, 600) = 16.46, $p < 0.001$). Dependent-sample *t*-tests were applied to compare head motion between every two groups. We found that that adults had significantly lower head motion relative to the full-term neonates ($t$ (368.44) = -8.67, $p < 0.001$), preterm neonates scanned at TEA ($t$ (79.69) = -4.14, $p < 0.001$), and preterm neonates scanned before TEA ($t$ (76.97) = -4.16, $p < 0.001$) (Figure S3). Bonferroni correction for multiple comparison was applied to statistical results.

*Figure S3 about here please*

***High-order networks functional connectivity in adults.*** In adults, a 2 × 3 repeated measure ANOVA [*type of FC* (within-network, between-network) × *network* (DMN, DAN, ECN)] showed a significant main effect of type of FC ($F$ (1, 175) = 2323.00, $p < 0.001$), which was driven by higher overall connectivity for the within- relative to between-network ($t$ (175) = 48.11, $p < 0.001$). We also found a main effect of network ($F$ (1.93, 337.773) = 37.50, $p < 0.001$), that was driven by lower overall connectivity for the DMN relative to the DAN ($t$(175) = -8.72, $p < 0.001$) and ECN ($t$ (175) = -3.71, $p < 0.001$) and lower overall connectivity for the ECN relative to DAN ($t$ (175) = -5.10, $p < 0.001$). Finally, a significant interaction effect of type of FC by network ($F$ (2, 350) = 96.49, $p < 0.001$) was driven by a smaller within- vs between-network FC difference in ECN relative to DMN ($t$ (175) = -11.20, $p < 0.001$) and DAN ($t$ (175) = -13.03, $p < 0.001$). Paired-t tests showed significantly higher within- to between-network FC for each network (DMN: $t$ (175) = 30.69, $p < 0.001$; DAN: $t$ (175) = 42.00, $p < 0.001$; ECN: $t$ (175) = 27.40, $p < 0.001$) (Figure S4), confirming that each of the three networks was differentiated as a cohesive unit in adults. Bonferroni correction for multiple comparison was applied to statistical results.

*Figure S4 about here please*

***The reciprocal relationship between the DMN and prefrontal networks in adults.*** A one-way ANOVA with repeated measures for between-network FC (DMN−DAN, DMN−ECN, DAN−ECN) showed a significant main effect ($F$ (1.93, 337.22) = 118.92, $p < 0.001$), which was driven by significantly lower FC in the DMN−DAN relative to the DMN−ECN ($t$ (175) = -6.19, $p < 0.001$) and DAN−ECN pairings ($t$ (175) = -14.01, $p < 0.001$), and significantly lower FC in the DMN−ECN relative to DAN−ECN pairing ($t$ (175) = -9.61, $p < 0.001$) (Figure S5). Bonferroni correction for multiple comparison was applied to statistical results. The lower FC between the DMN and DAN relative to the other pairings suggested a reciprocal relationship between the two networks in adults.

*Figure S5 about here please*

***Comparison of DMN-frontoparietal functional connectivity in neonates and adults.*** To investigate between-network connectivity in neonates relative to adults, we created a GLM that controlled for head motion, and compared each neonate group to the adult group. For full-term neonates, we found significant main effects of group for all of the pairings (DMN−DAN: $F$ (1, 455) = 278.77, $p < 0.001$; DMN−ECN: $F$ (1, 455) = 195.83, $p < 0.001$; DAN−ECN: $F$ (1, 455) = 15.59, $p < 0.001$), which was driven by significantly lower FC in DMN−DAN and DMN−ECN, and higher FC in DAN−ECN in the adults related to full-term neonates (Figure 7a). These results suggested that the DMN was more functionally differentiated from the DAN and ECN, and thus, suggesting a stronger reciprocal relationship in adults relative to full-term neonates. For preterm neonates scanned at TEA, we found a significant main effect of group for DMN−DAN ($F$ (1, 246) = 105.83, $p < 0.001$), DMN−ECN ($F$ (1, 246) = 91.39, $p < 0.001$), which was driven by significantly lower FC in

DMN−DAN and DMN−ECN in the adults related to preterm neonates scanned at TEA (Figure 7b). Similarly, to full-term neonates, these results demonstrated that the DMN was more functionally differentiated from DAN and ECN, suggesting a stronger reciprocal relationship in adults relative to preterm neonates scanned at TEA. We also compared between-network connectivity in preterm neonates scanned before TEA and adults using a GLM that controlled for head motion, although we did not observe a reciprocal relationship between DMN and frontoparietal network in that neonate group. We found a significant main effect of group for DMN−DAN ($F$ (1, 246) = 257.78, $p < 0.001$) and DMN−ECN ($F$ (1, 246) = 45.54, $p < 0.001$), which was driven by significantly lower FC in DMN−DAN and DMN−ECN in the adults relative to the preterm neonates scanned before TEA (Figure 7c).

### Supplementary Figures

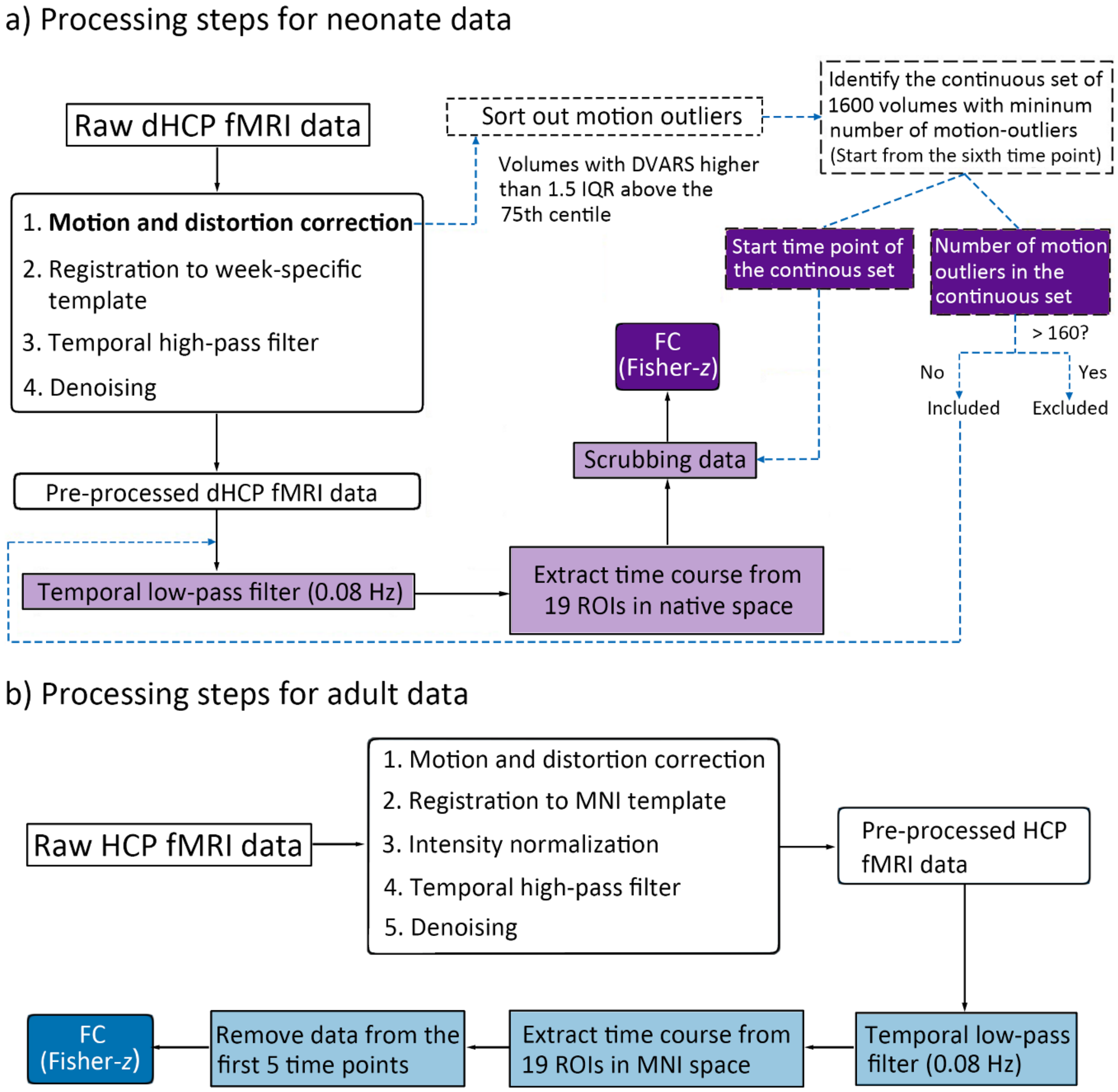


**Figure S1.** Data processing steps for neonate and adult data. a) Processing steps for the neonate rs-fMRI data. The rectangles with rounded corners indicate the steps in the fMRI neonatal pre-processing pipeline of Developing Human Connectome Project. The frames in light purple indicate additional processing steps in this study, and the dotted black line rectangles represent the steps for correcting motion outliers. The frames in dark purple represent data that we obtained. b) Processing steps for the adult fMRI data. The rectangles with rounded corners indicate the steps in the fMRI pre-processing pipeline of Human Connectome Project. The frames in light blue indicate additional processing steps in this study and the frame in dark blue represents data that we obtained. Abbreviations: dHCP, developing Human Connectome Project; DVARS, D referring to temporal derivative of time courses and VARS referring to root mean squared variance over voxels; ROI, regions of interest; IQR, Inter Quartile Range; FC, functional connectivity; HCP, Human Connectome Project.

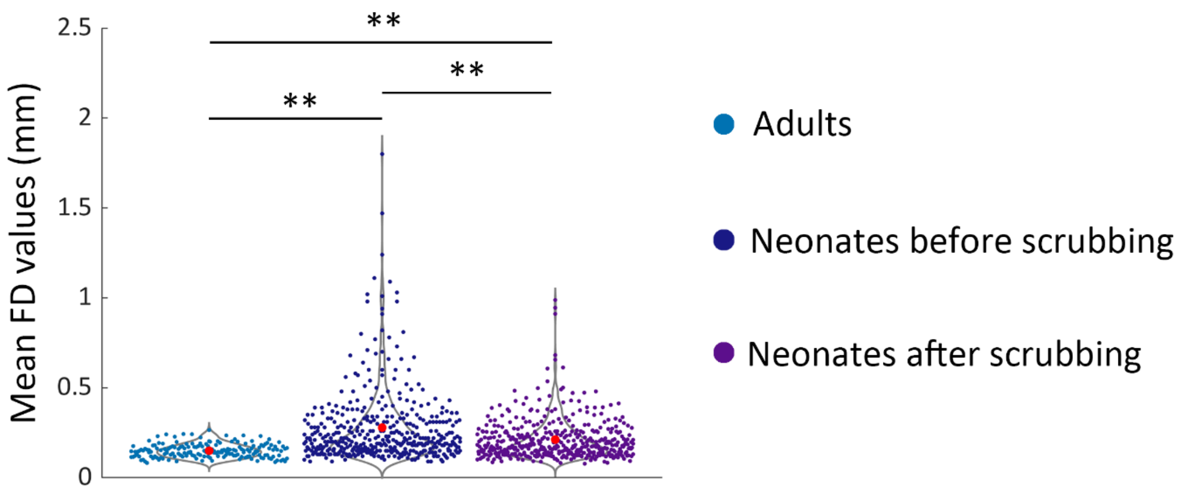


**Figure S2.** Head motion in neonates and adults. The red dot indicates mean framewise displacement value in each group. Abbreviations: FD, framewise displacement; ** = $p <$ 0.005.

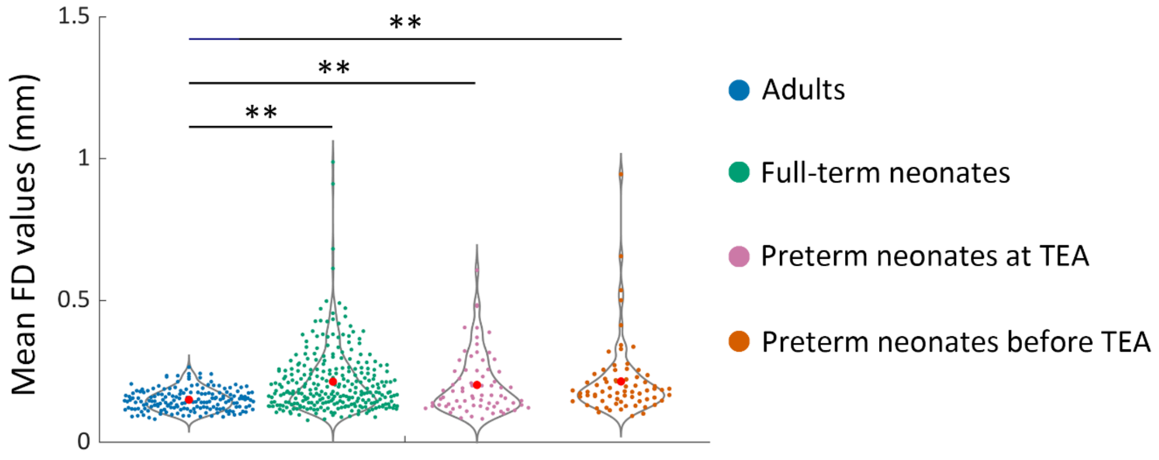


**Figure S3.** Head motion in the neonate groups after censoring and the adults. The red dot indicates mean framewise displacement value in each group. Abbreviations: FD, framewise displacement; TEA, term-equivalent age; ** = $p < 0.005$.

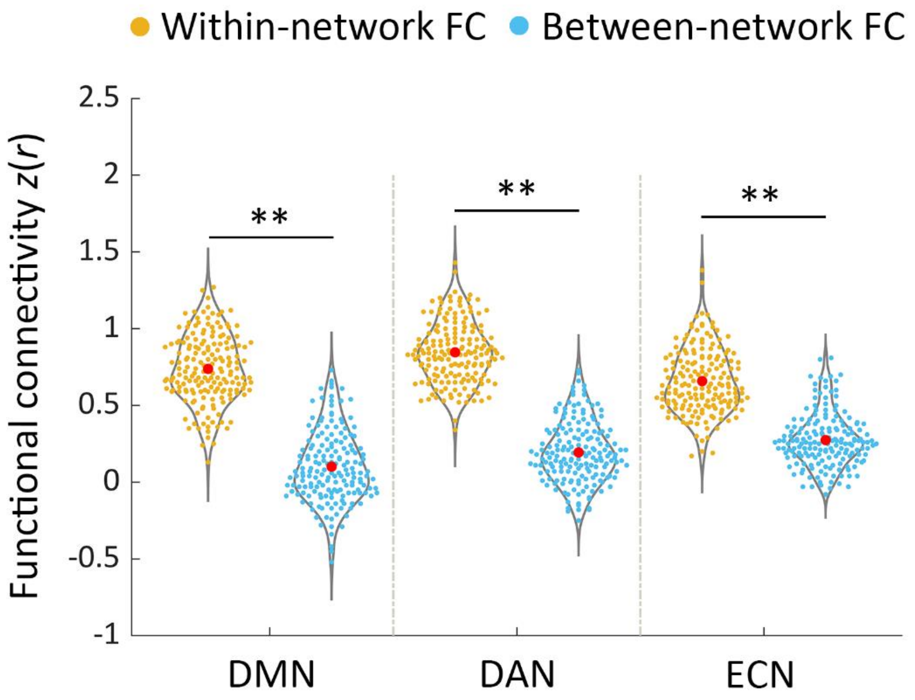


**Figure S4.** Network functional connectivity in adults. The between-network connectivity depicts the average FC of each network with the other two. The red dot indicates mean within/between-network functional connectivity of each network. Abbreviations: DMN, default mode network; DAN, dorsal attention network; ECN, executive control network; FC, functional connectivity; ** = $p < 0.005$.

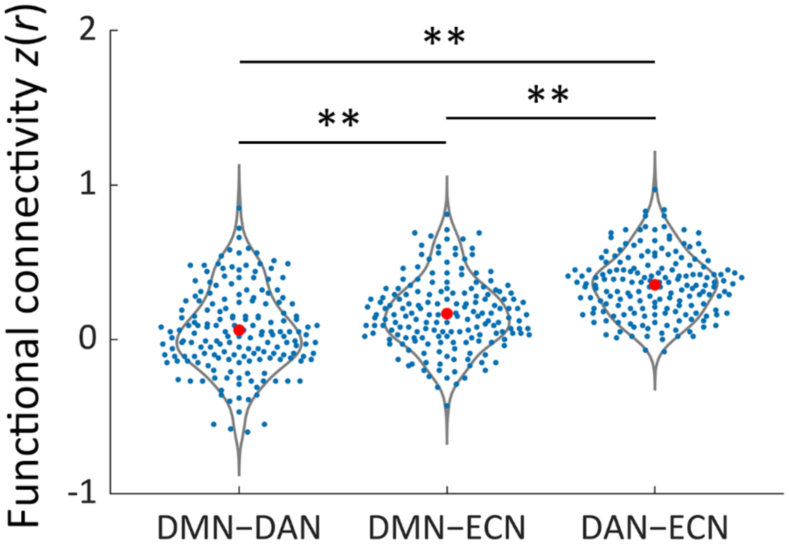


**Figure S5.** Between-network functional connectivity (FC) in adults. FC values were Fisher-*z* transformed. The red dot indicates mean functional connectivity in each network-pairing. Abbreviations: DMN−DAN, FC between the default mode network and dorsal attention network; DMN−ECN, FC between the default mode network and executive control network; DAN−ECN, FC between the dorsal attention network and executive control network; ** = $p < 0.005$.

### Supplementary Tables

**Table S1.** Previous rs-MRI studies of brain network development in neonates

| No. | Article | Subjects | GA at birth | Age at scan | State at scan | Scanner | TR (ms) | voxel size ($mm^3$) | Duration (min) | Head motion control | Template | Analysis method | Neural network identified |
|---|---|---|---|---|---|---|---|---|---|---|---|---|---|
| 1 | Fransson et al., (2007) | 12 preterm neonates scanned at TEA | 25 weeks 6 days (24 weeks 4 days –27 weeks 5 days) | 41 weeks 0 days PMA (39 weeks 1 day – 44 weeks 3 days) | under sedation | 1.5 T | 2000 | 2.8 × 2.8 × 4.5 | 10 | scrubbing | neonate brain template (Dehaene-Lambertz et al., 2002) | ICA | 1) VIS; 2) SMN; 3) AUD; 4) a network including the precuneus area, lateral parietal cortex, and the cerebellum; 5) an anterior network that incorporated the medial and dorsolateral prefrontal cortex. |
| 2 | Fransson et al., (2009) | 19 full-term neonates | 38 weeks 4 days (37 weeks 4 days – 39 weeks 1 day) | 40 weeks 2 days PMA (39 weeks 2 days – 41 weeks 6 days) | natural sleep | 1.5 T | 2000 | 2.8 × 2.8 × 4.5 | 10 | scrubbing | neonate brain template (Dehaene-Lambertz et al., 2002) | ICA | 1) VIS; 2) SMN; 3) bilateral temporal/inferior parietal cortex including the primary auditory cortex, 4)posterior lateral and midline aspects of the parietal cortex; 5) medial and lateral parts of the prefrontal cortex and 6) the bilateral basal ganglia. |
| 3 | Lin et al., (2008) | 38 preterm and full-term neonates | 35 – 42 weeks | 2 – 4 weeks after birth | natural sleep | 3T | 2000 | 4 × 4 × 4 | 5 | scrubbing | data-specific template | SCA | SMN and VIS (exists as early as 2 weeks after birth) |

| | | | | | | | | | | | | | |
|---|---|---|---|---|---|---|---|---|---|---|---|---|---|
| | | 26 one-year-olds born prematurely or at full-term age | N/A | N/A | | | | | | | | | |
| | | 21 two-year-olds born prematurely or at full-term age | N/A | N/A | | | | | | | | | |
| 4 | Doria et al., (2010) | 17 early preterm neonates scanned before TEA | 25 weeks 2 days – 31 weeks 1 day | 29 weeks 0 days – 32 weeks 1 day PMA | natural sleep | 3T | 1500 | 2.5 × 2.5 × 3.25 | 6.5 | FD > 5 mm | data-specific template | ICA and SCA | 1) In full-term neonates and preterm neonates scanned at TEA: VIS, AUD, SMN, motor, DMN, FPN, and ECN were completely present;<br>2) In early preterm neonates: AUD, dorsal visual stream and ECN were not found;<br>3) Before TEA, the DMN was incomplete. |
| | | 21 preterm neonates scanned before TEA | 26 – 35 weeks | 33 weeks 0 days – 36 weeks 4 days PMA | natural sleep (17/23) /under sedation | | | | | | | | |
| | | 24 preterm neonates scanned at TEA | 24 weeks 3 days – 35 weeks 3 days | 39 weeks 3 days – 43 weeks 2 days PMA | under sedation | | | | | | | | |
| | | 8 full-term neonates | 38 – 41 weeks 4 days | 39 weeks 1 day – 43 weeks 4 days PMA | natural sleep (6/8) /under sedation | | | | | | | | |
| 5 | Smyser et al., (2010) | 53 preterm neonates | 23 weeks 2 days – 34 weeks 0 day | 1) < 30 weeks PMA<br>2) 30 weeks PMA<br>3) 34.0 weeks PMA<br>4) 38.0 weeks PMA | natural sleep/resting quietly | 3T | 2910 | 2.4 × 2.4 × 2.4 | 10 | scrubbing | Adult template | SCA | 1) In preterm neonates and full-term neonates: motor-leg, motor-hand, motor-face, PCC, ACC, occipital, MPFC, LPFC, temporal, thalamus and cerebellum.<br>2) DMN precursor in full-term neonates. |
| | | 10 full-term newborns | N/A | Within 2—3 days of birth | | | | | | | | | |

| | | | | | | | | | | | | | |
|---|---|---|---|---|---|---|---|---|---|---|---|---|---|
| 6 | Alcauter et al., (2014) | 112 preterm and full-term neonates | 35 – 42 weeks | 4 weeks 5 days ± 2 weeks 5 days after birth | natural sleep | 3T | 2000 | 4 × 4 × 4 | 5 | scrubbing | data-specific template | SCA | 1) Neonates: Adultlike SMN, AUD, medial visual, and occipital pole and SN; Incomplete lateral visual, DMN, and FPN. 2) The lateral VIS, DMN, FPN showed dramatic synchronization during the first year and had minor refinement during the second year. |
| | | 129 one-year-olds | N/A | 1 year 32 days ± 35 days | | | | | | | | | |
| | | 92 two-year-olds | N/A | 2 year 32 days ± 33 days | | | | | | | | | |
| 7 | Gao et al., (2009) | 20 preterm and full-term neonates | 35 – 42 weeks | 3 weeks 3 days ± 1 week 5 days after birth | natural sleep | 3T | 2000 | 4 × 4 × 4 | 5 | scrubbing | data-specific template | ICA | An incomplete DMN is present in 2-week-olds. More adultlike DMN were found in 1-year-olds and 2-year-olds. |
| | | 24 one-year-olds | N/A | 1 year 1 month ± 1 month | | | | | | | | | |
| | | 27 two-year-olds | N/A | 2 years 1 month ± 1 month | | | | | | | | | |
| 8 | Gao et al., 2013 | 51 full-term and preterm neonates | 35 – 42 weeks | 3 weeks 2 days ± 1 week 5 days after birth | natural sleep | 3T | 2000 | 5 × 4 × 4 | 5 | None | a subject not included in this study and scanned at 3 weeks | SCA | 1) Neonates: incomplete DMN and DAN. 2) 1-year-olds: highly synchronized DMN and DAN. |
| | | 50 one-year-olds | N/A | 1 year 1 month ± 1 month | | | | | | | | | |
| | | 46 two-year-olds | N/A | 2 years ± 1 months | | | | | | | | | |

| | | | | | | | | | | | | | |
|---|---|---|---|---|---|---|---|---|---|---|---|---|---|
| 9 | Gao et al., (2015a) | 65 full-term neonates | 35 – 42 weeks | < 1 month (N = 45)<br>3 months (N = 34)<br>6 months (N = 33)<br>9 months (N = 29)<br>12 month (N = 35) | natural sleep | 3T | 2000 | 4 × 4 × 4 | 5 | scrubbing | data-specific template and adult MNI template | SCA | 1) Neonates: SMN, AUD, VN; incomplete lateral visual/parietal network, SN, DMN, FPN.<br>2) 1-year-olds: adultlike lateral visual/parietal network and DMN; incomplete SN and FPN |
| 10 | Gao et al., (2015b) | 143 full-term neonates | 36 – 42 weeks | 4 weeks 5 days ± 2 weeks 5 days after birth (N = 112)<br><br>1 year old (N = 129)<br><br>2 years old (N = 92) | natural sleep | 3T | 2000 | 4 × 4 × 4 | 5 | scrubbing | data-specific template | ICA | 1) Neonates: VN and SMN;<br>2) The auditory/language, lateral visual/parietal, DMN, right FPN, SN, and left FPN were topologically incomplete and isolated in neonates but demonstrated consistent synchronization during the first 2 years of life. |
| 11 | Wylie et al., (2014) | 12 full-term neonates | N/A | 8 weeks 0 days ± 2 weeks 6 days after birth | under sedation | 3T | 2000 | 3.43 × 3.43 × 3.4 | N/A | motion larger than 1 voxel | MNI template | ICA | DMN, VIS, AUD, SMN, basal ganglia, precuneus, visual spatial, language, ECN, anterior SN. |
| 12 | Damaraju et al., 2014 | 4-month-old full-term infants<br><br>9-month-old full-term infants | N/A | 18 weeks 4 days ± 2 weeks 1 day after birth<br><br>40 weeks 4 days ± 1 week 2 days after birth | natural sleep | 3T | 2000 | thickness = 3.5 mm | 8.3 | despiking step | 9-month MNI template (Altaye et al., 2008) | ICA | In both groups: sub-cortical, AUD, VIS, SMN, DMN, temporal, attentional and frontal network, cerebellum. |

| | | | | | | | | | | | | | |
|---|---|---|---|---|---|---|---|---|---|---|---|---|---|
| 13 | He et al., (2015) | 27 preterm neonates scanned at TEA | 26 weeks 6 days ± 2 weeks 0 days | 39 weeks 4 days ± 1 week 3 days PMA | natural sleep | 3 T | 3000 | slice thickness = 3 mm | 5.2 | scrubbing | neonate brain template (Kuklisova-Murgasova et al., 2011) | ICA | FPN, ECN, motor, SMN, medial visual, occipital visual and lateral visual areas. |
| 14 | He et al., (2016) | 34 preterm neonates | ≤ 32 weeks | 32 weeks 4 days ± 1 week 0 days PMA (N = 19)<br><br>39 weeks 2 days ± 1 week 2 days PMA (N = 22)<br><br>52 weeks 6 days ± 1 week 4 days PMA (N = 25) | natural sleep | 3 T | 3000 | slice thickness = 3 mm | 5.2 | scrubbing | neonate brain template (Kuklisova-Murgasova et al., 2011) | ICA | In all the three groups: The occipital visual, medial visual, lateral visual, AUD, motor, SMN, cerebellum, brainstem, DMN, ECN and FPN. |
| 15 | Ball et al., (2016) | 105 preterm neonates scanned at TEA<br>26 full-term neonates | 30 (23 – 34) weeks<br><br>39 (37 – 41) weeks | 42 (39 – 48) weeks PMA<br><br>43 (39 – 46) weeks PMA | under sedation | 3 T | 1500 | 2.5 × 2.5 × 4 | 6.4 | ICA + FIX clean up | preterm brain template (Serag et al., 2012) | ICA | For both groups: frontal, parietal, temporal, occipital cortex, basal ganglia and cerebellum. |
| 16 | Weinstein et al., (2016) | 32 preterm neonates | 29 weeks 0 days ± 2 weeks 5 days | 37 weeks 4 days ± 1 week 4 days PMA | natural sleep | 3 T | 3000 | slice thickness = 3 mm | N/A | None | None | SCA | Motor and VIS |
| 17 | Cui et al., (2017) | 44 preterm neonates scanned before TEA | 24 weeks 5 days – 32 weeks 2 days | 32 weeks 1 day ± 1 week 5 days PMA (29 weeks 6 days – 35 weeks 4 days) | N/A | 3 T | 2000 | 4 × 4 × 4 | N/A | None | Montreal Neurological Institute pediatric atlas | ICA | Medial visual, lateral visual, AUD, SN, motor, prefrontal network, brainstem and thalami, frontal cortical network and cerebellum. |

| | | | | | | | | | | | | | |
|---|---|---|---|---|---|---|---|---|---|---|---|---|---|
| 18 | Linke et al., (2018) | 11 preterm neonates (No neuropathology) | 27 weeks 4 days (25 – 34) | 37 weeks 0 days PMA (35 – 42) | natural sleep with playing lullabies | 1.5 T | 1920 | 3 mm isotropic resolution | 7 | average >2 mm in two or more of the four fMRI runs | UNC neonatal brain template (Shi et al., 2011) | ICA | For both preterm scanned at TEA and full-term neonates: motor, AUD, VIS, ECN and DMN. |
| | | 19 preterm neonates (Neuropathology) | 27 weeks 4 days (24 – 36) | 37 weeks 4 days PMA (35 – 41) | | | | | | | | | |
| | | 3 full-term neonates (No neuropathology) | 40 weeks 0 days (39 – 41) | 40.5 (40 – 41) weeks PMA | | | | | | | | | |
| | | 7 full-term neonates (Neuropathology) | 40 weeks 0 days (38 – 41) | 41.0 (39 – 43) weeks PMA | | | | | | | | | |
| 19 | Rajasilta et al., (2020) | 21 full-term neonates | 39 weeks 6 days ± 1 week 1 day | 3 weeks 5 days ± 1 week 0 days after birth | natural sleep | 3T | 2500 | 3 × 3 × 3 | 6 | motion larger than 3 mm in multiple time points | UNC2 neonate T2 template (Shi et al., 2011) | ICA | VIS, AUD, thalamic, basal ganglia, cerebellar and brainstem, insular, SMN, motor, DMN, prefrontal, frontal, parietal and temporoparietal networks. |

GA , gestational age; PMA, postmenstrual age; ICA, Independent Component Analysis; SCA, Seed Correlation Analysis; FD, framewise displacement; TEA, term-equivalent age; SMN, sensorimotor network; VIS, visual network; DMN, default mode network ; AUD, auditory network; FPN, frontoparietal network; ECN, executive network; SN, salience Network; ACC, anterior cingulate cortex; PCC, posterior cingulate cortex; MPFC, medial prefrontal cortex; LPFC, lateral prefrontal cortex

**Table S2.** Information obtained from the developing Human Connectome Project for scans used in the present study.

| **Group** | **Sex** | **Birth age (GA, weeks ± days)** | **Scan age (PMA, weeks ± days)** | **Birth weight (Kg)** |
|---|---|---|---|---|
| Full-term neonates | 160M/122F | 40.0 ± 8.6 | 41.2 ± 12.0 | 3.35 ± 0.54 |
| Preterm neonates scanned at TEA | 41M/32F | 32.0 ± 25.6 | 40.9 ± 14.5 | 1.76 ± 0.79 |
| Preterm neonates scanned before TEA | 50M/23F | 32.5 ± 13.4 | 34.6 ± 13.4 | 1.78 ± 0.61 |

Abbreviations: GA, gestational age; PMA, postmenstrual age; TEA, term-equivalent age; M, male; F, female.

**Table S3.** Regions of interests for the default mode network, dorsal attention network and executive control network (from Raichle 2011).

| Network | Index | ROI | MNI coordinates | | |
|---|---|---|---|---|---|
| | | | *x* | *y* | *z* |
| **Default Mode Network** | 1 | Posterior cingulate/precuneus | 0 | -52 | 27 |
| | 2 | Medial prefrontal | -1 | 54 | 27 |
| | 3 | Left lateral parietal | -46 | -66 | 30 |
| | 4 | Right lateral parietal | 49 | -63 | 33 |
| | 5 | Left inferior temporal | -61 | -24 | -9 |
| | 6 | Right inferior temporal | 58 | -24 | -9 |
| **Dorsal Attention Network** | 7 | Left frontal eye field | -29 | -9 | 54 |
| | 8 | Right frontal eye field | 29 | -9 | 54 |
| | 9 | Left posterior IPS | -26 | -66 | 48 |
| | 10 | Right posterior IPS | 26 | -66 | 48 |
| | 11 | Left anterior IPS | -44 | -39 | 45 |
| | 12 | Right anterior IPS | 41 | -39 | 45 |
| | 13 | Left MT | -50 | -66 | -6 |
| | 14 | Right MT | 53 | -63 | -6 |
| **Executive Control Network** | 15 | Dorsal medial PFC | 0 | 24 | 46 |
| | 16 | Left anterior PFC | -44 | 45 | 0 |
| | 17 | Right anterior PFC | 44 | 45 | 0 |
| | 18 | Left superior parietal | -50 | -51 | 45 |
| | 19 | Right superior parietal | 50 | -51 | 45 |

IPS: Intraparietal sulcus, MT: Middle temporal area, PFC: Prefrontal cortex
